# Closed-loop optimization of a high-dimensional generative latent space for rhythmic visual response

**DOI:** 10.64898/2026.06.27.734819

**Authors:** Jesse A. Livezey, Yaqing Su, Stephanie Wolfer, Abigaïl Ingster, David J. Klein, Adam Hanina

## Abstract

Neural oscillations accompany a wide range of cognitive states and behaviors including perception, memory, and movement, and modulating them is of growing interest for both basic neuroscience and clinical research. Previous demonstrations of closed-loop modulation of visual neural responses mainly relied on invasive recordings and focused on firing-rate maximization rather than rhythmic modulation. Here, we show that the relative power of steady-state visual evoked response (SSVEP), measured with readily-available, non-invasive electroencephalography, can be modulated in closedloop as a function of image stimulus parameters for single participants within a single session. Stimulus optimization with flicker frequencies in the alpha and theta bands was successful in 10, 20, and 40 dimensional latent spaces. We also show that optimized stimuli generalize to new participants when shown in open-loop. Finally, we characterize the visual features that modulate relative SSVEP power and find that low-frequency spatial power in the image drives theta and alpha in opposite directions. Together, our results show that closed-loop stimulus optimization is a viable method for rhythmic neural modulation using noninvasive neuroimaging methods.

## 1 Introduction

Rhythmic ensemble neural activity (neural oscillations) has been found to robustly associate with perceptual processing, behaviors and mental states. In visual processing, the alpha (8-14 Hz) oscillation has been shown to phase-lock to saccades during visual scene encoding [1], and visual attention in both human and macaque monkeys is modulated by theta (4-7 Hz) [2, 3]. In speech processing, delta (1-4 Hz), alpha, and theta neural oscillations, measurable with EEG, track both acoustic and linguistic features in the speech signal [4]. Beta oscillations (13-30 Hz) and their coupling are shown to be involved in motor preparation [5, 6] and maintenance-of-state [7]. Meanwhile, abnormalities in neural oscillations are shown to correlate with dysfunctions in sensory, motor, and cognitive processes (see review by [8]), leading to a growing body of research on invasive and noninvasive interventions via the modulation of oscillatory activities. For example, neuromodulation of gamma-band activity using visual stimulation has been indicated as a potential tool for slowing down cognitive decline in older subjects and patients with Alzheimer’s disease [9, 10, 11]. Non-invasive electrical stimulation has been suggested as a method for partially restoring vision with activity in the alpha band as a specific neuromodulatory target [12]. However, existing techniques for modulating oscillations are highly constrained in their parameter space, for example, by only changing the amplitude (intensity) of the stimulation, thus cannot be easily optimized or adapted to individuals, resulting in high cross-subject differences in their outcomes [13, 14, 15, 16].

One promising approach for modulating neural oscillations in individual subjects is closed-loop neural optimization. To achieve this, a predefined “neural objective” in response to the input stimulus at trial *t* is fed back to a stimulus synthesizer to alter the parameter for the input at trial *t* + 1, in order to alter the neural feature in a designated direction, e.g., maximizing the amplitude of alpha-band activity at a certain electrode. Such a realtime closed-loop approach has been successfully utilized in visual [17, 18, 19] and motor [20, 21] systems in animal models. However, application in human subjects remain limited largely due to the poor signal-to-noise ratio in noninvasive measurements especially from relatively low-cost devices such as electroencephalography (EEG) [22, 23]. Some existing studies of closed-loop EEG optimization [24, 25] require the participant’s active but implicit effort, making the optimization procedure long and results less interpretable and generalizable [26]. Moreover, previous stimulus optimization studies often operate in a very low dimensional space where it is possible to test a regular grid of parameters, which is not feasible in even 10s of dimensions. For example, combinations of color or luminance and frequency within the gamma band for entrainment [27] or shape size and contrast for modulation of endogenous theta[28]. Even when generative models were used for stimulus generation, the stimuli produced vary along simple dimensions of grayscale luminance and basic shapes [23].

In this study, we establish the feasibility of a closed-loop visual stimulus optimization pipeline using widely available EEG hardware and passive video viewing, readily exploitable for both fundamental and clinical research. We implemented and validated a closed-loop system that optimizes human rhythmic EEG response to visual textures parametrized in the latent space of a deep generative image model. The generative image model we use can generate high dimensional (up to ∼16, 000) naturalistic color images and textures, although here we parametrically constrain the search space to a subspace of the full dimensionality to understand the dependence of optimization effect on dimensionality. Our neural objective is based on steady-state visual evoked potentials (SSVEP) features, a common technique used in visual neuroscience [29, 30, 31, 32] and braincomputer interface research [33, 34, 35] to boost the signal-to-noise ratio of EEG-based features. We show that stimulus parameters that maximize and minimize our neural objective can be identified within a single 30-minute session in a 10-dimensional search space. We then show that the same system is also able to optimize in 20- and 40-dimensional spaces, although the optimization effect size, which we quantify with Hedges’ *g* (an unbiased version of the more familiar Cohen’s *d* [36, 37]), is smaller for larger dimensionalities. At the individual participant level, participant-specific factors explain more of the observed variability in optimization effect size than dimensionality or neural band. Next, we demonstrate that the stimulus latent parameters and extracted low-level visual features can both be used to train encoding models to predict neural objectives in new participants. The distribution of power in spatial frequencies is the most consistent predictor of neural objectives with low spatial frequencies having opposite effects on theta and alpha bands. Finally, we visualize trajectories in stimulus space that maximally modulate the neural objective for all dimensionalities and both neural bands. Together, these results show that closed-loop optimization in humans using EEG is a viable approach for producing stimuli optimized to drive neural targets of neuroscientific and neuromodulatory interest.

## 2 Results

A total of 30 subjects participated in two types of electroencephalography (EEG) experiments without overlapping: *optimization* (*n* = 24) and *generalization* (*n* = 6). In both types of experiments, we defined the “neural objective” as the log ratio of the spectral power density (PSD) of the EEG signal at theta (6 Hz) or alpha (10 Hz) flicker frequency to the adjacent frequency bins averaged across 10 occipital and occipital-parietal electrodes in the same 2 second window as the stimulus presentation. In *optimization* experiments, subjects were instructed to passively watch short flickering videos while their real-time EEG response was used to generate new videos for optimizing, that is, maximizing or minimizing, their neural objectives using our closed-loop optimization system. Videos generated in optimization experiments were pooled and filtered for the *generalization* experiments, where participants were similarly instructed to passively watch short video clips, but the videos were predefined and did not adapt to their EEG response.

In *optimization* experiments, three different dimensionalities of stimulus parameter (texture) space were tested at each flicker frequency: 10, 20, and 40. In *generalization* experiments, all three dimensionalities were tested at 10 Hz whereas only 40-dimensional stimuli were tested at 6 Hz. We refer to each combination of experiment type, flicker frequency, and stimulus parameter dimension, as an experiment *version*. For *optimization* experiments, each participant was tested for two separate experiment runs, each consisting of a single version without repetition. All experiment versions were run in 6 participant except for Theta 40d which was run in 12 participants although only the first 6 participants were used for generalization. For *generalization* experiments, all versions of the same flicker frequency were aggregated in the same experiment run. Each experiment version lasted approximately 30 minutes or less.

### 2.1 Closed-loop stimulus optimization setup

Each optimization experiment was initiated with 25 “preset” trials which were used to warm-start optimization and common to all participants within an experiment version. The remaining 225 trials were a mix of 150 “maximization” trials, 50 “minimization” trials, and 25 trials repeating the preset trials. The 225 trials were evenly split into 25-trial blocks and the within-block order was pseudorandomized per-participant, maintaining a roughly constant fraction of each type in every 25 trial block (Fig. 7**A**). Note that the fraction denotes the average number of trials in each block, for example a total of 225*/*9 = 45 preset trials are uniformly distributed among the 26th to 250th trials, but the exact number of preset trials varies between blocks since 45 does not divide evenly into 25. Trials consist of 2 seconds of fixation, 2 seconds of stimulation, and 2 seconds of rest (Fig. 7**B**). The optimization space consisted of a softmax-weighted linear combination of 10, 20, or 40 latent textures. Fig. 7**C** shows these latent textures projected onto image space with the top row, two rows, and all rows being used for 10, 20, and 40d optimization, respectively. The same 10, 20 and 40d textures were collected for both flicker frequencies. In addition, a second Alpha 10d set of experiments were collected, referred to as “Alpha2 10d” with a different set of 10 textures outlined in red. Extended data Figure 13 shows which participants ran which experiment versions. See Methods for more details on the experiment design.

**Figure 1:**
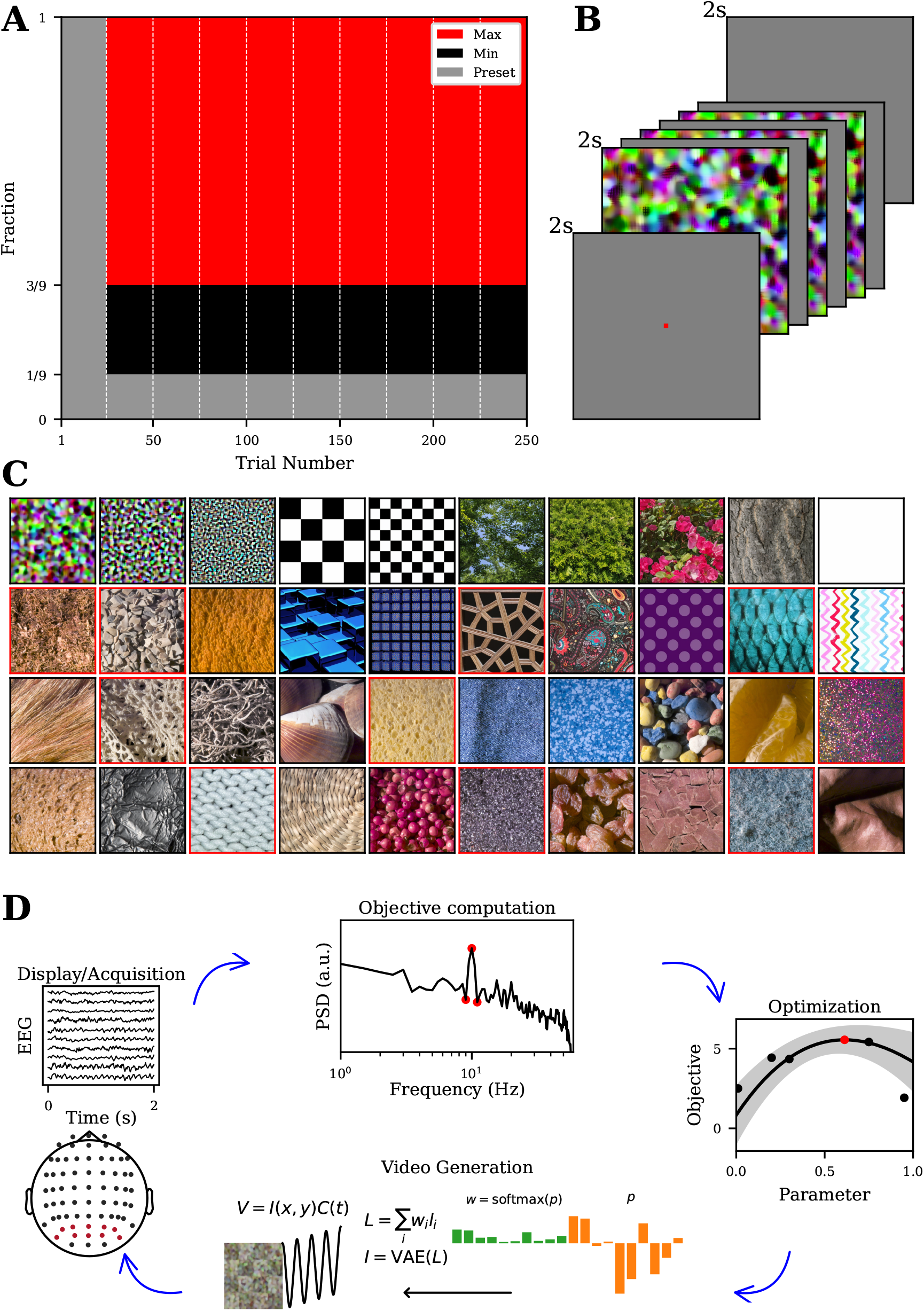
Experimental structure for closed-loop stimulus optimization. **A:** Block and trial structure for a single optimization experiment. Vertical dashed lines indicate blocks of 25 trials. The vertical axis indicate the fraction of trials of preset (gray), minimize (black), and maximize (red) types in each block. **B:** Each trial is composed of 2 seconds of a gray screen with a red fixation point, 2 seconds of stimulation with a red fixation dot, and 2 seconds of a gray screen. **C:** Images shown are generated from single elements of the 40 latent textures that define the optimization spaces. Experiments with *d* = 10 use latent textures from the top row with the exception of the Alpha2 experiment which uses the textures outlined in red, *d* = 20 use latent textures from the top 2 rows, and *d* = 40 use all latent textures. **D:** Diagram of the closed-loop experimental structure starting from the bottom and proceeding clockwise is as follows: video generation from stimulus parameters, video display and raw EEG acquisition, objective computation from EEG, optimization of the stimulus parameters for the next trial. EEG electrodes used for neural objective calculation are highlighted in red.

**Figure 2:**
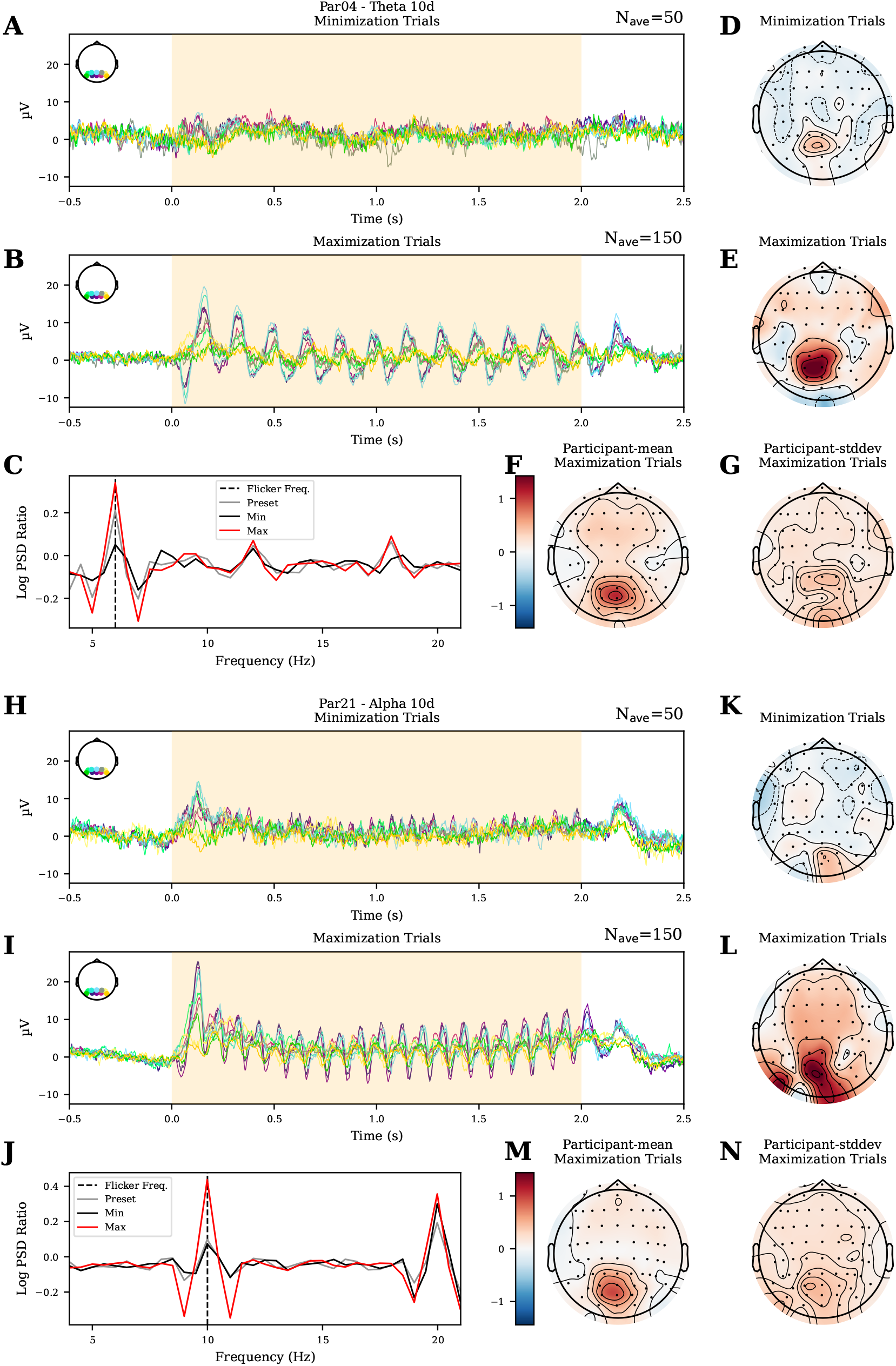
Visual stimulation robustly evokes SSVEP for Theta and Alpha 10d. **A-B:** Median evoked potentials per trial type (minimization and maximization) for the 10 occipital channels used for optimization in the participant with the largest optimization effect for the Theta 10d experiment (Par04). EEG signals were common-average referenced and bandpass filtered between 1 and 50 Hz for visualization. **C:** Mean log power spectral density (PSD) ratios are shown for each trial type for the participant from **A**. Dashed black line indicates the flicker frequency. **D-E:** The log PSD ratio is calculated per electrode and the mean across all trials is plotted for minimization and maximization trials for the participant from **A**. Colormap is symmetrical around zero with blue indicating values less than zero and red indicating values greater than zero. **F-G:** The mean and standard deviation across participants of the per-electrode mean log PSD ratio is plotted for maximization trials. **H-N:** The same plots are shown for Alpha 10d where Par21 is used in **H-L**. (difference in mean is 0.37, Welch’s t-test, *p <* 1e-26, df = 82.4). The log power spectral density ratio is strongest over occipital electrodes, although this participant shows a higher degree of lateral asymmetry than the Theta 10d participant (Fig. 9K-L). The cross-participant mean and standard deviations of the means both show structure in occipital electrodes (Fig. 9M,N). Together, these results demonstrate that the flicker-induced SSVEP is spectrally and spatially localized for both Theta and Alpha 10d, and that SSVEP amplitude varies across different optimization trial types in a controlled fashion.

**Figure 3:**
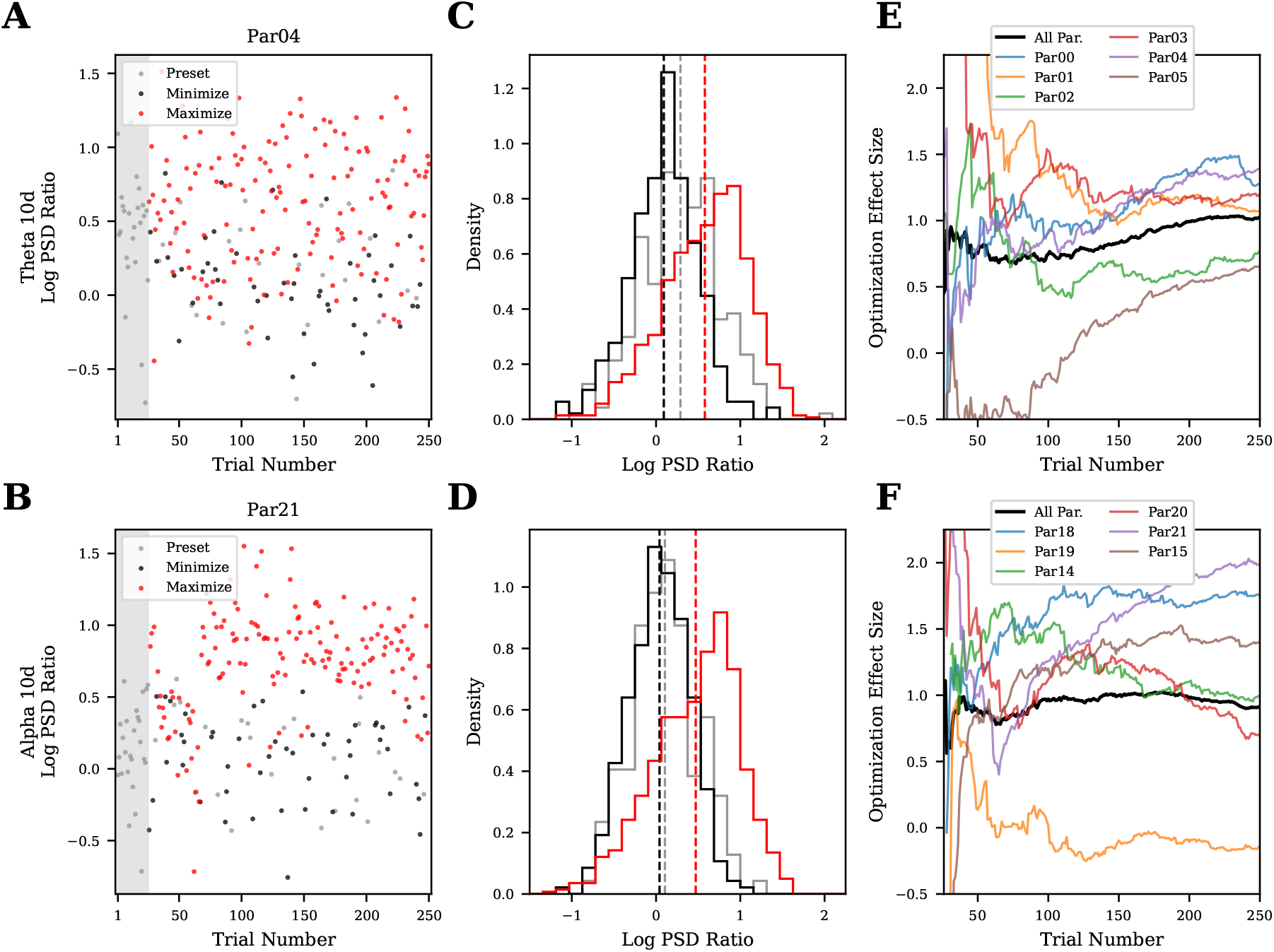
Visual stimulation can be reliably optimized in a single session for Theta and Alpha 10d. **A**,**B:** For participant 04 (Par04) in 10d Theta and participant 21 (Par21) in 10d Alpha, the per-trial neural objective is shown for preset (gray), minimize (black) and maximize (red) trials. The gray rectangle indicates the first block which is entirely composed of preset trials. **C**,**D:** Neural objective histograms across all participants for each flicker frequency. Vertical dashed lines indicates means. Colors as in **A**,**B. E**,**F:** Hedges’ g is plotted against trial number where trials up-to and including the trial number are used. Colored lines are individual participants and the black line combines trials across all participants.

**Figure 4:**
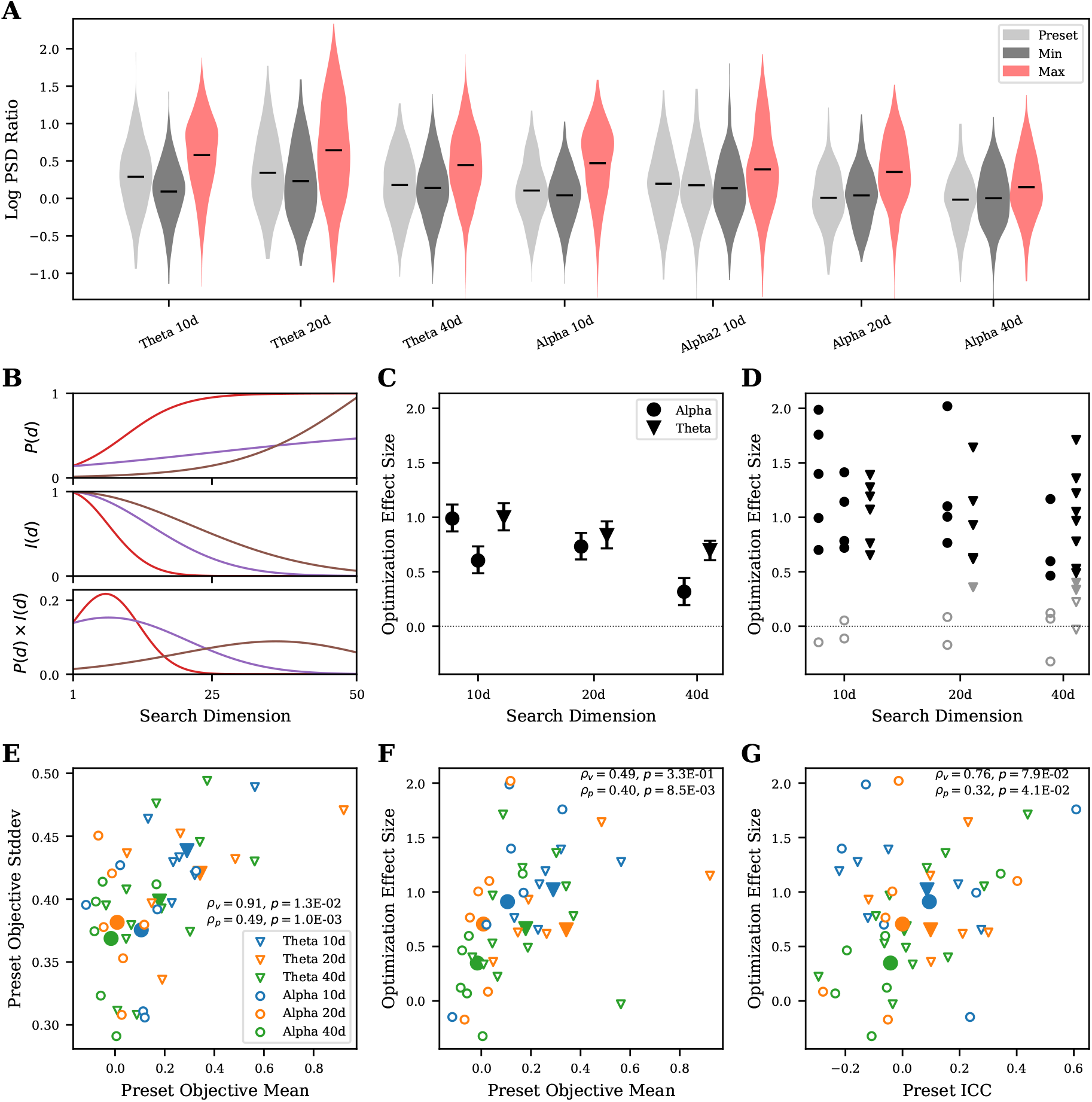
Optimization effect size depends on experiment version and individual participant. **A:** For each optimization experiment version, the Log SSVEP Ratio for each trial type (color) and search dimension (horizontal-axis group) is summarized in a violin plot. The means are indicated with horizontal lines. **B:** Three models of how generative search may scale with search dimension. Each color represents a different optimization problem. In the top row, a model for how infinite-data performance (*P* (*d*)) scales with search dimension. The second row shows a model of how the information gained in a single trial (*I*(*d*)) scales with search dimension. The third row models the optimization experiment performance as the product of *P* (*d*) and *I*(*d*). **C:** The Optimization Effect Size is shown for Alpha and Theta datasets combining trials across all participants. Errorbars indicate 95% bootstrapped confidence intervals. In **C, D**, the Alpha 10d points are ordered by Alpha then Alpha2 from left to right. **D:** The Optimization Effect Size is shown for Alpha and Theta datasets for all dimensions and participants. False discovery rate corrected p-values are indicated by marker color and style. Solid black points indicates *p <* 0.01, solid gray indicates 0.05 *> p* ≥ 0.01, and outlined gray indicates *p* ≥ 0.05. Shapes as in **C. E:** For the preset trials, the mean and standard deviation of the Log SSVEP Ratio are scattered for each participant (small unfilled markers). For each experiment version, the average mean and standard deviation across participants is shown (large solid markers). Shape indicates band and color indicates dimensionality. The linear correlation coefficient and p-value are shown for the within-experiment version (*ρ*_*v*_) and within-participant (*ρ*_*p*_) points. **F:** The preset objective mean and optimization effect size are scattered for each participant. Shape and color as in **E**. The linear correlation coefficient and p-value is shown. **G:** ICC calculated on the preset trials is scattered against optimization effect size for individual participants (small unfilled markers, ICC(1,1) and across participants (large solid markers, ICC(C,1)). The linear correlation coefficient and p-value is shown for the within-experiment version (*ρ*_*v*_) and within-participant (*ρ*_*p*_) points Shape and color as in **E**.

**Figure 5:**
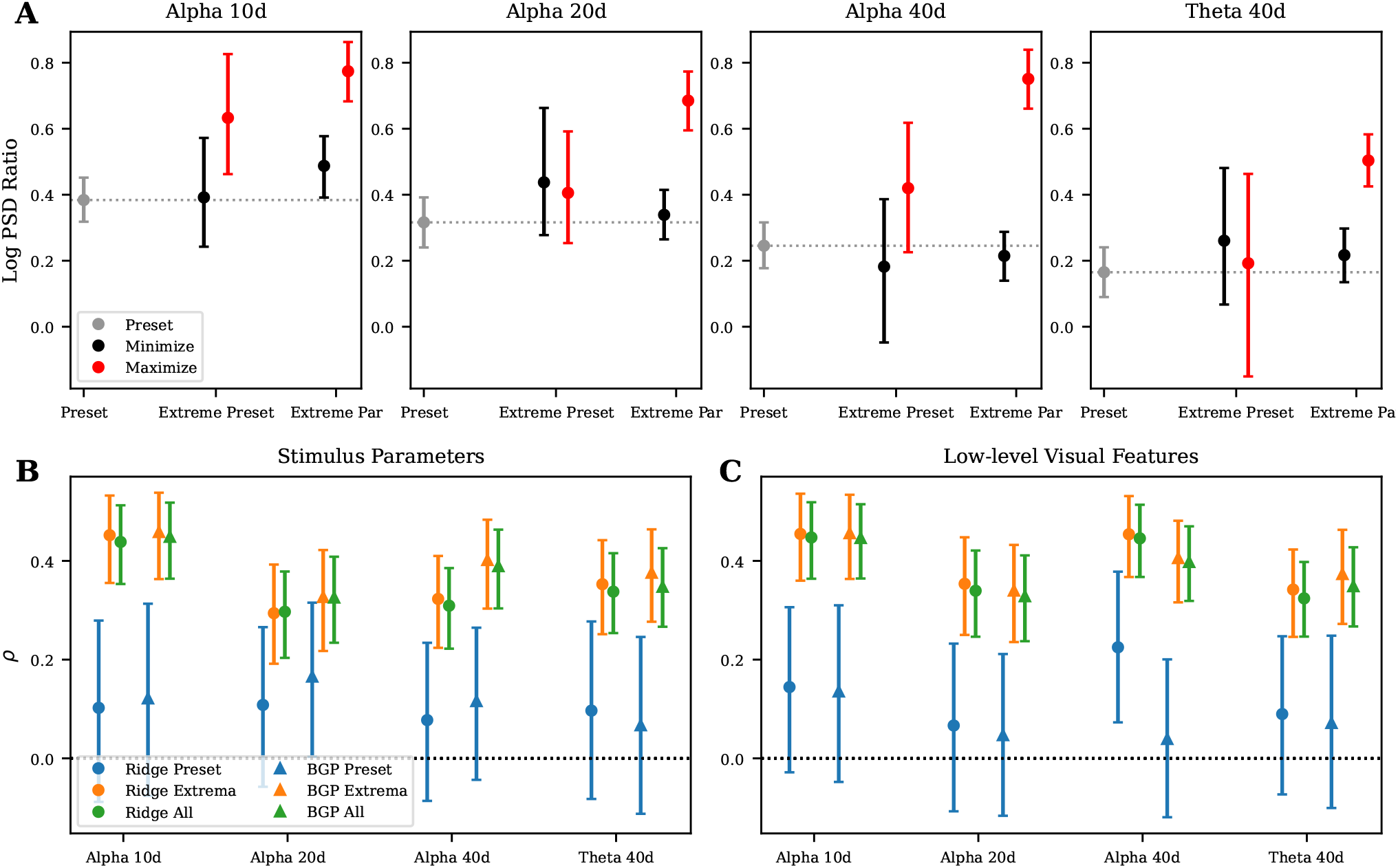
Optimized stimuli generalize to new participants. **A:** Mean and 95% bootstrapped confidence intervals across generalization participants are shown for neural objectives collected from Preset stimuli, preset stimuli with the min/max average neural objective in optimization participants (Extreme Preset), and stimuli with the min/max neural objective from each optimization experiment (Extreme Par) Stimuli from all Alpha dimensions and Theta 40d were collected and are shown across panels. **B:** Mean correlations and 95% bootstrapped confidence intervals on the generalization trials are shown for preset stimuli (blue markers), extreme stimuli (orange markers) and all stimuli (green markers) for crossvalidated Ridge (circles) and Bayesian Gaussian process (BGP) regression (triangles) models trained on the optimization datasets to predict neural objectives from stimulus parameters. **C:** Mean correlations and 95% bootstrapped confidence intervals on the generalization trials are shown for models trained to predict neural objectives from a set of 9 low-level image features computed from the generated texture. Colors and shapes as in **B**.

**Figure 6:**
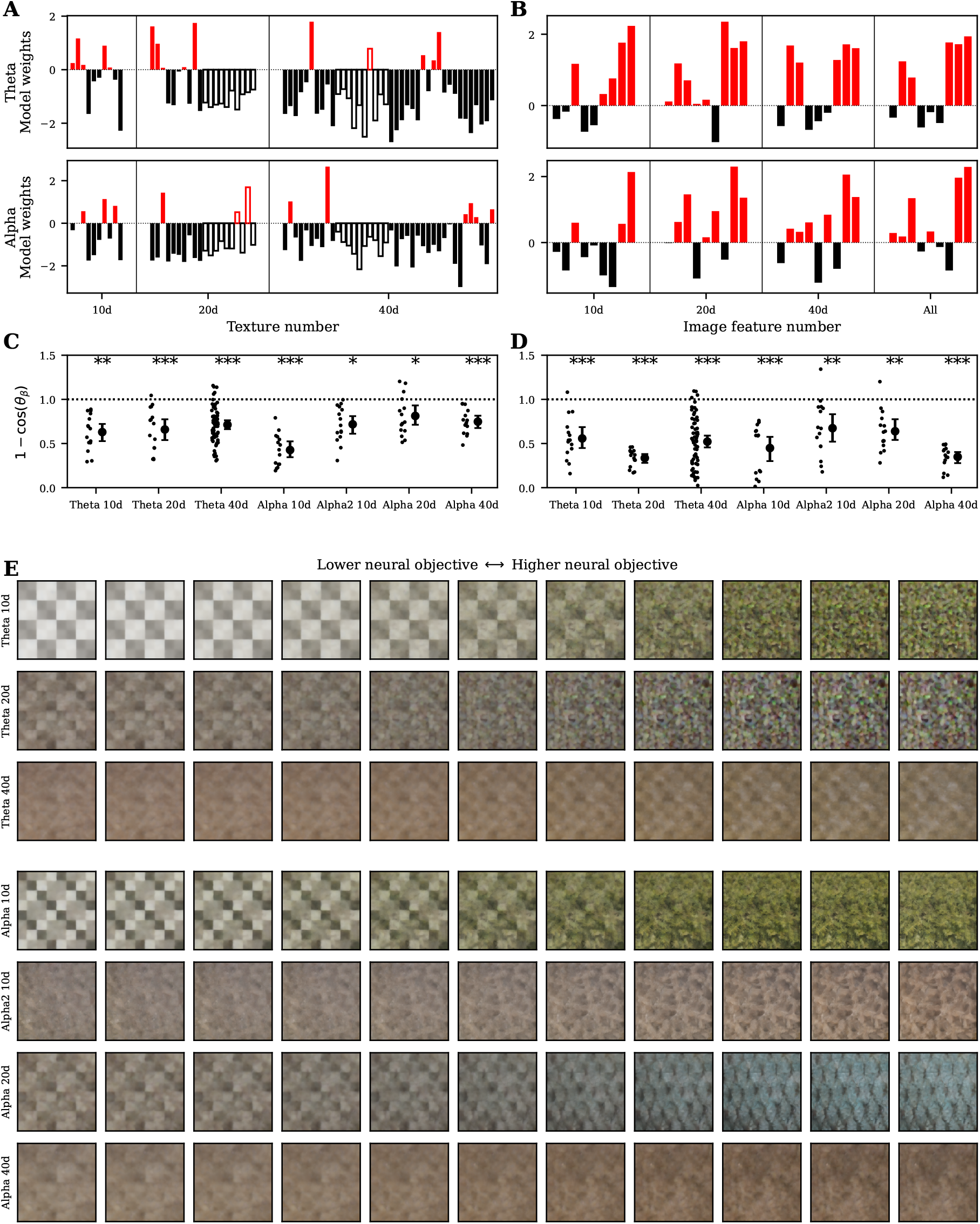
Modulation axes are reliable across participants in lower dimensions. **A:** Ridge regression weights (normalized by their standard deviation) for models trained on 10, 20, and 40d (left to right) and Theta (top) and Alpha (bottom) experiment versions to predict neural objectives from stimulus parameters. The first 10 weights are filled, the second 10 weights are empty bars, and the last 20 weights are filled bars. Red or black bars indicate a positive or negative weights. **B:** Ridge regression weights normalized by their standard deviation for models trained on all Theta (top) and Alpha (bottom) experiment versions for models trained to predict neural objectives from a set of 9 features computed from the generated texture. Colors as in **A. C:** Pairwise cosine distances between participant’s Ridge weights train on stimulus parameters (small dots) and the mean distance and bootstrapped 95% confidence intervals (large dots with errorbars) are shown for all experiment versions. Dashed horizontal line at 1 is the mean of the null distribution assuming randomly distributed model weights. **D:** Pairwise cosine distances between participant’s Ridge weights trained on low-level image features are shown for all experiment versions. Markers, null line, and p-values as in **C. E:** A set of textures spanning the linear encoding subspace of a Ridge model is shown for each experiment version.

**Figure 7:**
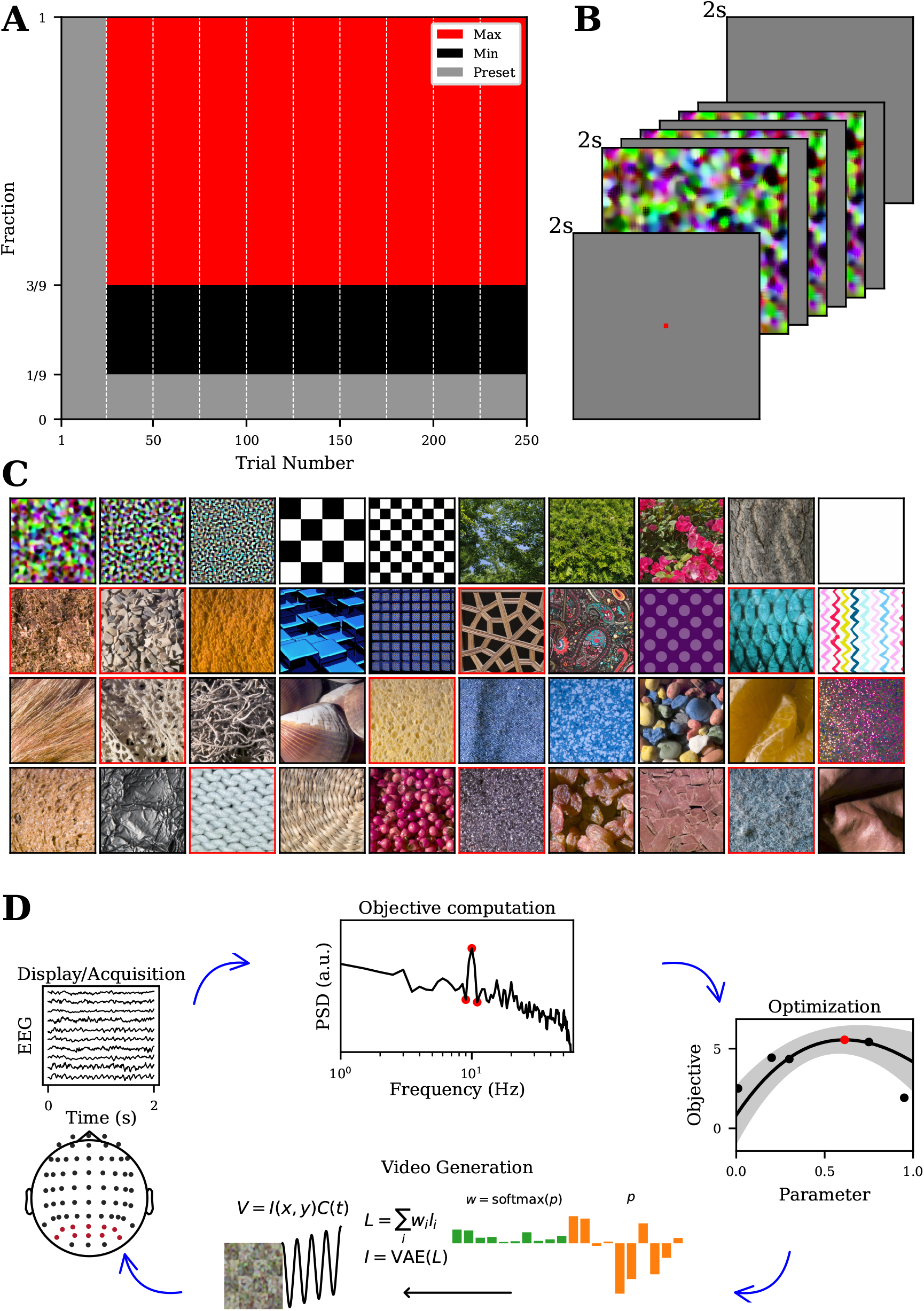
Experimental structure for closed-loop stimulus optimization. **A:** Block and trial structure for a single optimization experiment. Vertical dashed lines indicate blocks of 25 trials. The vertical axis indicate the fraction of trials of preset (gray), minimize (black), and maximize (red) types in each block. **B:** Each trial is composed of 2 seconds of a gray screen with a red fixation point, 2 seconds of stimulation with a red fixation dot, and 2 seconds of a gray screen. **C:** Images shown are generated from single elements of the 40 latent textures that define the optimization spaces. Experiments with *d* = 10 use latent textures from the top row with the exception of the Alpha2 experiment which uses the textures outlined in red, *d* = 20 use latent textures from the top 2 rows, and *d* = 40 use all latent textures. **D:** Diagram of the closed-loop experimental structure starting from the bottom and proceeding clockwise is as follows: video generation from stimulus parameters, video display and raw EEG acquisition, objective computation from EEG, optimization of the stimulus parameters for the next trial. EEG electrodes used for neural objective calculation are highlighted in red.

Figure 7D shows a diagram of the closed-loop experimental structure. Starting at the bottom and proceeding in a clockwise direction: first, stimulus parameters, *p*, are converted to weights, *w* by a softmax operation. These weights are used to form a linear combination of the latent textures in Figure 7**C**. The latent texture is passed through the decoder of a variational autoencoder (VAE) taken from a latent diffusion model [38, 39] to create an image, and the image pixels are modulated sinusoidally, at the given flicker frequency, between the image and a uniform gray background to produce a video. Second, the video is displayed to the participant while the raw EEG signal is streamed through Lab Stream Layer (LSL) [40]. Third, the raw EEG undergoes minimal online preprocessing and the log power spectral density ratio neural objective is calculated for the occipital and occipital-parietal channels shown in red in the topoplot (see EEG preprocessing and neural objective calculation for details). Finally, the paired stimulus parameters and objective are added to the trial history and a Bayesian optimizer is used to select the next set of parameters to test. This process is repeated throughout the duration of the experiment.

### 2.2 Visual stimulation robustly evokes SSVEP for Theta and Alpha 10d

Periodic flickering of visual stimulus features is a commonly used technique to evoke steady-state visual evoked potentials (SSVEP) [29], which manifest as sharp peaks in neural power spectral densities at the flicker frequency and its harmonics. Low-level visual stimulus features have been found to modulate the amplitude and phase of SSVEP [30]. Here, we first sought to show that our stimulus optimization space could be used to controllably and differentially evoke SSVEP. For minimization and maximization trials, the optimized videos were able to differentially evoke SSVEP amplitude in occipital electrode. Figure 8A and B show the median SSVEP for minimization trials and maximization trials, respectively, for the participant that had the highest optimization effect in the Theta 10d experiment (Par04). The maximization trials show entrainment to the 6 Hz flicker stimulation starting just after stimulus onset (*t* = 0) and subsiding one cycle after stimulus offset (*t* = 2).

**Figure 8:**
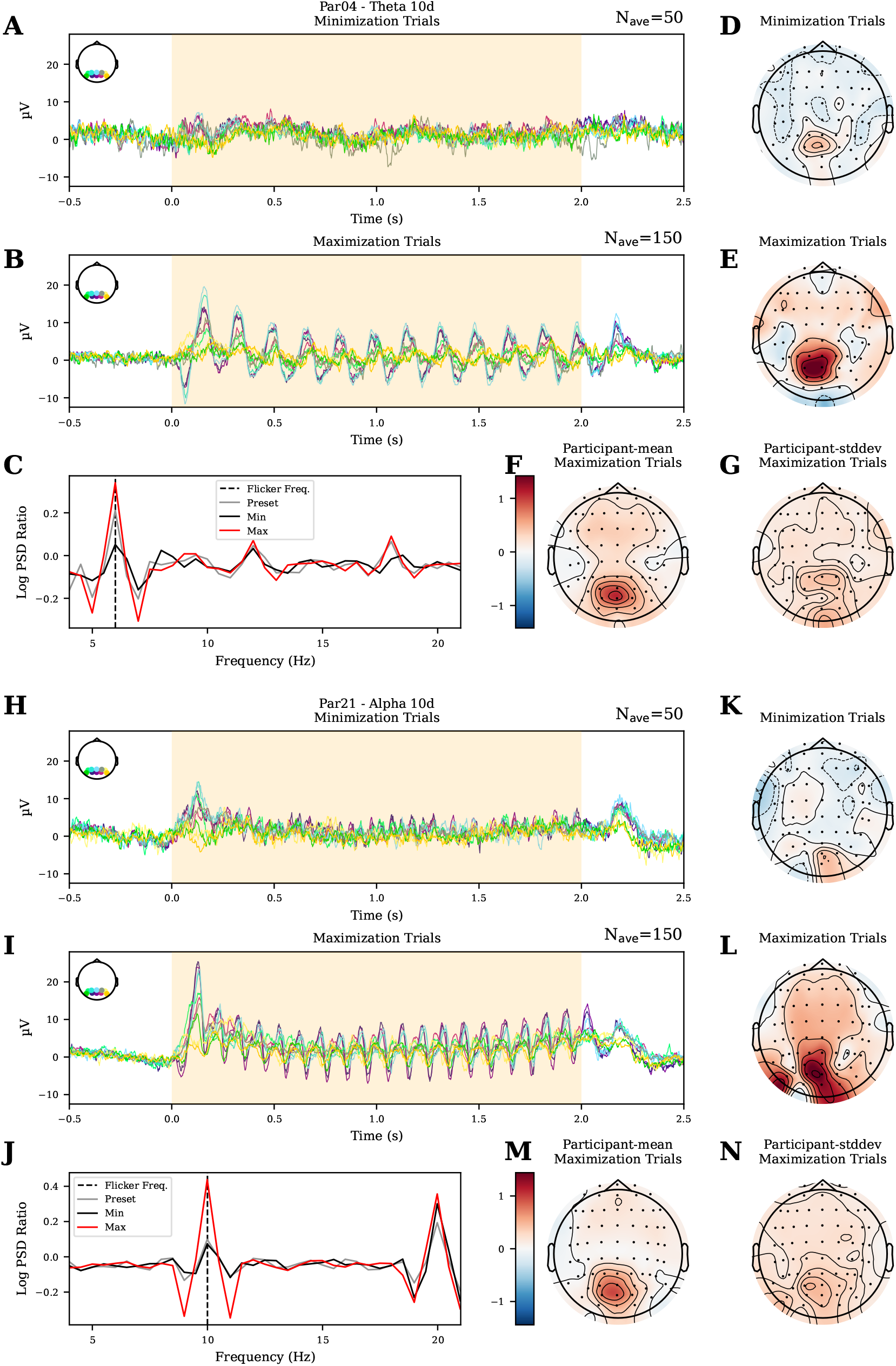
Visual stimulation robustly evokes SSVEP for Theta and Alpha 10d. **A-B:** Median evoked potentials per trial type (minimization and maximization) for the 10 occipital channels used for optimization in the participant with the largest optimization effect for the Theta 10d experiment (Par04). EEG signals were common-average referenced and bandpass filtered between 1 and 50 Hz for visualization. **C:** Mean log power spectral density (PSD) ratios are shown for each trial type for the participant from **A**. Dashed black line indicates the flicker frequency. **D-E:** The log PSD ratio is calculated per electrode and the mean across all trials is plotted for minimization and maximization trials for the participant from **A**. Colormap is symmetrical around zero with blue indicating values less than zero and red indicating values greater than zero. **F-G:** The mean and standard deviation across participants of the per-electrode mean log PSD ratio is plotted for maximization trials. **H-N:** The same plots are shown for Alpha 10d where Par21 is used in **H-L**.

We next looked to characterize the spectral and spatial characteristics of the SSVEP signal. We found that the SSVEP is spectrally localized to the flicker frequency and its harmonics with small amounts of evoked power leaking into neighboring frequency bins. Figure 8C plots the log of the power spectral density ratio between the flicker frequency and the average of the 2 frequency bins *±* 2 bins away around the flicker frequency, averaged across trials and EEG electrodes used for computing the neural objective. The stimulation period was 2 seconds, therefore the frequency resolution is limited to 0.5 Hz. The maximization trials show a significantly larger mean log power spectral density ratio compared to the minimization trials at the flicker frequency (difference in mean is 0.29, Welch’s t-test, *p <* 1e-17, df = 93.0). Figure 8D,E show the mean per-electrode log power spectral density ratio taken over minimization and maximization trials, respectively. Spatially, high log power spectral density ratio values are clustered over occipital electrodes, with smaller overall amplitudes in minimization trials compared to maximization trials. Across participants, the mean and standard deviation of the neural objective for maximization trials mainly peaked around occipital electrodes (Fig. 8F,G).

Qualitatively, the SSVEP looks similar for the participant (Par21) that had the highest optimization effect size for Alpha 10d (Fig. 9H-N). The maximization trials show a significantly larger mean log power spectral density ratio compared to the minimization trials at the flicker frequency

**Figure 9:**
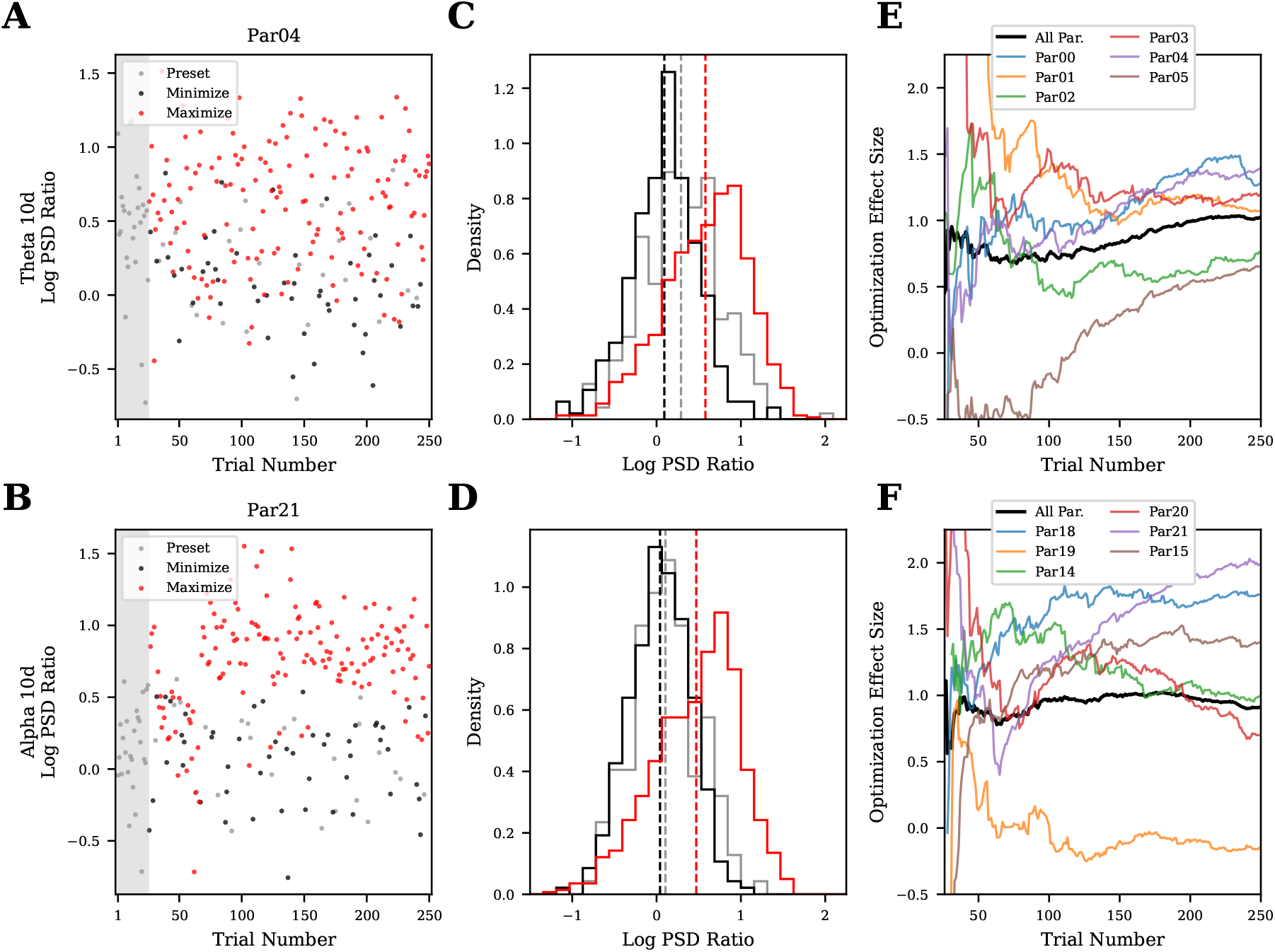
Visual stimulation can be reliably optimized in a single session for Theta and Alpha 10d. **A**,**B:** For participant 04 (Par04) in 10d Theta and participant 21 (Par21) in 10d Alpha, the per-trial neural objective is shown for preset (gray), minimize (black) and maximize (red) trials. The gray rectangle indicates the first block which is entirely composed of preset trials. **C**,**D:** Neural objective histograms across all participants for each flicker frequency. Vertical dashed lines indicates means. Colors as in **A**,**B. E**,**F:** Hedges’ g is plotted against trial number where trials up-to and including the trial number are used. Colored lines are individual participants and the black line combines trials across all participants.

### 2.3 SSVEP is reliably optimized for Theta and Alpha 10d

For optimization to be considered successful, it must be able to drive the distribution of evoked neural objectives apart between minimization versus maximization trials: the farther apart the distributions, the more effective the optimization is. Here, we quantify the optimization effect size by Hedges’ *g*, which is derived from Cohen’s *d* with a bias adjustment for small sample sizes [37].

Figure 9A,B show neural objectives recorded in each trial for the participants with highest optimization effect sizes for Theta and Alpha 10d, participants Par04 and Par21 respectively. Visually, objectives in maximization and minimization separate after about 75 trials in both participants. To quantify the separation across all participants, we aggregate and histogram all trials. Figure 9C,D show histograms of the neural objectives for preset (gray), minimization (black), and maximization trials (red) for trials taken from all participants for Theta and Alpha, respectively. The mean neural objective in the maximization trials is significantly higher than those in the minimization trials (difference in means is 0.49 for Theta 10d *p <* 1e-52, df = 619.6; difference in means is 0.43 for Alpha 10d *p <* 1e-49, df = 721.7). For both flicker frequencies the mean preset neural objective falls between the minimization and maximization trials. This shows that optimized stimuli are able to modulate the distribution of neural objectives between minimization and maximization trials for Theta and Alpha 10d.

In general, Bayesian optimization methods trade-off so-called exploration and exploitation at a trial-by-trial basis, testing places where the surrogate model (see Optimizer and regression models for more detail) has high uncertainty as quantified by the predicted standard deviation (“exploration”) and places where the surrogate model predicts good objective values as quantified by the predicted mean (“exploitation”). In particular, we use the upper confidence bound acquisition function to adjudicate between candidate parameters, which selects points that have high (or low for minimization) predicted means and large standard deviations. This tradeoff and the single-trial variability of the neural recordings mean that the per-trial optimization effect size is not expected to strictly monotonically increase. To quantify how optimization unfolds across trials, we calculate the cumulative optimization effect size at each trial number, taking all minimization and maximization trials up to that point into account (Fig. 9E,F). Each colored line represents an individual participant, and the black line combines trials from all participants. For Theta 10d, the effect size tends to increase across trials and starts to plateau around 225 trials for some participants. For Alpha 10d, 3 of the participants’ effect sizes generally increase over trials (Par18, Par15, Par21), 2 participants’ peak around the 100th trials (Par14, Par20). However, 1 participant’s effect size hovers around 0 (Par19), indicating unsuccessful optimization. Note the effect sizes estimated with small numbers of trials (near the left extreme of the plot) are likely unreliable since they are estimated from a small number of trials. By the final trial, all Theta 10d participants have significant optimization effect sizes greater than 0.5 (*p <* 1e-6, Bonferroni-corrected Welch’s t-test, *n* = 6) and 5 of 6 Alpha 10d participants have effect sizes greater then 0.5 (*p <* 1e-6, Bonferroni-corrected Welch’s t-test, *n* = 6) and 1 with an effect size near zero. Together, the aggregated and trial-wise optimization effect sizes both indicate that Theta and Alpha SSVEP can indeed be reliably optimized in a 10d latent texture space.

### 2.4 Optimization effect size depends on experiment version and individual participant

Higher-dimensional stimulus parameter search spaces would allow more varied visual statistics to be searched over. Successful optimization in higher dimensional stimulus parameter spaces would enable a more detailed understanding of visual function and stronger and more targeted neuromodulatory effects. However, blackbox optimization will only scale so far before the exponential growth of the search space as a function of dimensionality makes optimization exponentially more difficult: the “curse of dimensionality” [41]. It is not currently known how closed-loop, blackbox optimization will scale as a function of parameter space dimensionality in humans with EEG-based neural objectives.

In order to determine how optimization scales with stimulus parameter dimensionality, we ran optimization experiments with a fixed trial budget in latent texture spaces of 10, 20, and 40 dimensions. In addition, we ran a second set of Alpha 10d experiments with a set of 10 non-overlapping latent textures (Alpha2 10d). Figure10A summarizes the distributions of neural objectives recorded in all versions of optimization experiments. Gray, black, and red violin plots show the distribution of preset, minimization, maximization trials, respectively and are grouped horizontally by experiment version. For the Alpha2 10d experiment version, 25 preset trials from the main Alpha 10d experiment were included in addition to the Alpha2 10d preset trials to aid in comparing optimization at fixed dimension across latent textures. The corresponding neural objectives are summarized separately in the left and right gray violin plots in the Alpha2 10d group, respectively. For all experiment versions, maximization trials have a significantly higher mean than minimization or preset trials (*p <* 1e-6, Welch’s t-test, Bonferroni-corrected with *n* = 14).

Conceptually, there are two main effects that determine the success of blackbox optimization as a function of parameter dimensionality. Figure 10B illustrates three hypothetical sets of scaling curves. As the search space grows, the optimal stimulus in the search space can only get closer to the globally optimal stimulus (or reach a plateau). This means that the potential for optimization effect size (*P* (*d*)) should monotonically increase as a function of dimensionality (Fig. 10B, top panel). The competing effect is that the amount of information gained per trial (*I*(*d*)) about the stimulus-response function goes down as the search space grows (Fig. 10B, middle panel). This is the curse of dimensionality. Together, these two effects mean that optimization effect size (proportional to the product *P* (*d*) *× I*(*d*), Fig. 10, B bottom panel) should increase to a peak at some dimensionality that depends on the specific shapes of these two effects, then decrease as the curse of dimensionality overtakes the benefit of increasing the stimulus parameter space dimensionality. In the red example, the growing stimulus space quickly saturates (*P* (*d*)) and the information gained per trial (*I*(*d*)) drops quickly, together leading to a sharp increase then decrease in optimizer effectiveness around 5d. In the brown example, *P* (*d*) does not saturate over the whole range and *I*(*d*) has a much slower drop, leading to an optimization effectiveness curve that increases up to 40d before starting to decrease.

**Figure 10:**
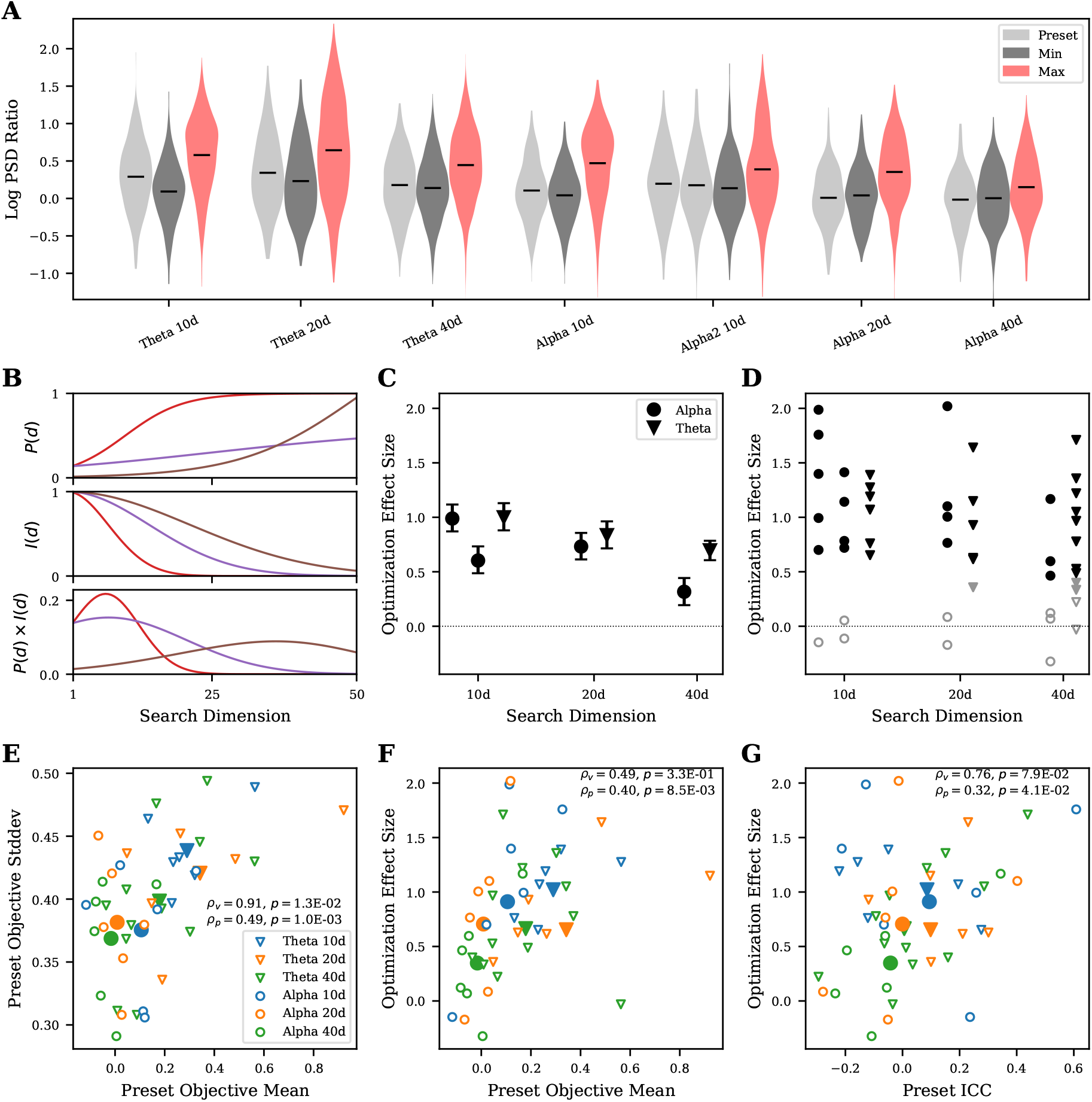
Optimization effect size depends on experiment version and individual participant. **A:** For each optimization experiment version, the Log SSVEP Ratio for each trial type (color) and search dimension (horizontal-axis group) is summarized in a violin plot. The means are indicated with horizontal lines. **B:** Three models of how generative search may scale with search dimension. Each color represents a different optimization problem. In the top row, a model for how infinite-data performance (*P* (*d*)) scales with search dimension. The second row shows a model of how the information gained in a single trial (*I*(*d*)) scales with search dimension. The third row models the optimization experiment performance as the product of *P* (*d*) and *I*(*d*). **C:** The Optimization Effect Size is shown for Alpha and Theta datasets combining trials across all participants. Errorbars indicate 95% bootstrapped confidence intervals. In **C, D**, the Alpha 10d points are ordered by Alpha then Alpha2 from left to right. **D:** The Optimization Effect Size is shown for Alpha and Theta datasets for all dimensions and participants. False discovery rate corrected p-values are indicated by marker color and style. Solid black points indicates *p <* 0.01, solid gray indicates 0.05 *> p* ≥ 0.01, and outlined gray indicates *p* ≥ 0.05. Shapes as in **C. E:** For the preset trials, the mean and standard deviation of the Log SSVEP Ratio are scattered for each participant (small unfilled markers). For each experiment version, the average mean and standard deviation across participants is shown (large solid markers). Shape indicates band and color indicates dimensionality. The linear correlation coefficient and p-value are shown for the within-experiment version (*ρ*_*v*_) and within-participant (*ρ*_*p*_) points. **F:** The preset objective mean and optimization effect size are scattered for each participant. Shape and color as in **E**. The linear correlation coefficient and p-value is shown. **G:** ICC calculated on the preset trials is scattered against optimization effect size for individual participants (small unfilled markers, ICC(1,1) and across participants (large solid markers, ICC(C,1)). The linear correlation coefficient and p-value is shown for the within-experiment version (*ρ*_*v*_) and within-participant (*ρ*_*p*_) points Shape and color as in **E**.

The purple example falls between these two. The details of each experimental design and participant will potentially have its own set of scaling curves which could depend on factors including how the search space manifold is constructed at each dimensionality, individual differences in structural or functional organization of participants’ brains, and the quality of the neural recording.

In order to determine how optimization scales with parameter space dimensionality in the paradigm presented here, we ran experiment versions with 10, 20, and 40 dimensional stimulus parameter spaces. For all experiment versions except for Alpha2 10d, the smaller stimulus parameter spaces are subspaces of the larger parameter spaces and are the same for theta and alpha bands (Figure 7). Figure 10C shows optimization effect sizes and 95% bootstrapped confidence intervals for all experiment versions, as quantified by Hedges’ *g* computed by combining all minimization and maximization trials across participants. Note that the left and right Alpha 10d markers are for Alpha 10d and Alpha2 10d, respectively. We find that all experiment versions have meaningful and statistically significant optimization effect sizes (*p <* 2e − 5, Bonferroni corrected t-tests, *n* = 7). Effect sizes range from 0.6 to 1.0 for all experiment versions except for Alpha 40d where it is 0.32. For the main set of Theta and Alpha experiment versions (excluding Alpha2 10d) the effect size decreases as a function of dimensionality: 10d: 1.00, 20d: 0.85, 40d: 0.70 and 10d: 0.99, 20d: 0.73, 40d: 0.32, respectively. This indicates that the main set of experiment versions are in the dimensionality regime where optimization can still be reliably successful, but increased dimensionality beyond 10d makes closed-loop optimization less effective.

For the main set of experiment versions, we had to make an arbitrary choice about which textures to start with in 10d and how to grow our search parameter space. This leaves the question of whether a different choice of how to structure the search parameter space could have led to a different scaling across dimensionalities. To partially answer this question, we created the Alpha2 10d experiment version, which uses 10 textures that are only in the Alpha 20d and 40d experiment versions. Alpha2 10d has an optimization effect size of 0.60 (Fig. 10C, Alpha 10d marker on the right), which is lower than both the Alpha 10d and 20d experiment versions. One possible explanation for this is that the latent textures chosen for the Alpha 10d experiment are closer to the optimal stimulus compared to those chosen for the Alpha2 10d experiment, which means that for the same experiment paradigm, specific choices such as how to grow the parameter search space can impact optimization effect size scaling by altering *P* (*d*) (Fig. 10B) in this case.

Unlike open-loop experiments where all participants can be shown the same set of stimuli, closedloop experiments adapt to individual participants. This can potentially lead to more varied results per-participant since optimization outcomes are sensitive to both cross-participant variability and and the cross-trial dynamics of Bayesian optimization. When broken-out by participant, we find that there is substantial cross-participant variability in optimization effect size. Figure 10D shows the per-participant optimization effect size across all experiment versions. Marker shape and horizontal-axis order is the same as in Figure 10C. Here, marker color and fill indicate Benjamini-Hochberg false discovery rate corrected p-values with filled black points indicating *p <* 0.01, filled gray points for 0.05 *> p* ≥0.01, and open gray points for *p* ≥0.05. The spread in per-participant optimization effect sizes is large compared to the confidence intervals estimated for the experiment version-level optimization effect sizes, indicating that participant-specific factors may explain some of the variability in each experiment version.

We next sought to explain this variability in optimization effect sizes across participants and experiment versions. For individual participants within an experiment version, the preset trials are a shared set of trials which can be used to estimate overall SSVEP susceptibility through the preset objective mean and how the cross-stimulus variance is partitioned between noise and structure through an intra-class correlation (ICC)[42]. Across participants, there is substantial variability in the distribution of neural objectives on the preset trials. Figure 10E shows the mean (horizontal axis) and standard deviation (vertical axis) for individual participants for each experiment version (small unfilled markers) and the means of these quantities across participants for each experiment version (large solid markers). The correlation across all participants and experiment versions (*ρ*_*p*_ = 0.49, *p* = 1.0e-3, *n* = 42) and for averages within experiment versions (*ρ*_*v*_ = 0.91, *p* = 1.3e-2, *n* = 6) are significant. Across participants and experiment versions, the preset trial neural objective mean is correlated with the optimization effect size (Fig. 10F, *ρ*_*p*_ = 0.40, *p* = 8.5e-3, *n* = 42) and across experiment versions there is a medium but statistically underpowered correlation (*ρ*_*v*_ = 0.49, *p* = 3.3e-1, *n* = 6). Figure 10G shows the optimization effect size scattered against the reliability of the preset objectives as quantified by ICC (1,1) for individual participants and ICC(C,1) for each experiment version. The correlation across all participants and experiment versions (*ρ*_*p*_ = 0.32, *p* = 4.1e-2, *n* = 42) is significant, and the correlation across experiment versions is large but statistically underpowered (*ρ*_*v*_ = 0.74, *p* = 9.4e-2, *n* = 6). In order to disentangle the impact of the participant and experiment version factors, we ran a Type 2 ANOVA to determine the effect that preset mean, ICC, log-parameter dimensionality, and binary categorical neural band have on optimization effect size. We found that there was an overall statistically significant effect (*F* = 3.83, *p* = 0.011, see A.2 for more details). While the variances explained by individual factors are generally not significant, we note that participant-specific factors have the largest sum-of-squares followed by the log-parameter dimensionality. The neural band has a negligible sum-of-squares. Together, this shows that participant-specific factors explain the majority of optimization effect size variability at the participant level.

### 2.5 Optimized stimuli generalize to new participants

Although the optimization effect size shows a high degree of variability across individual participants, it is not clear to what degree neural response functions are aligned across participants since the generated videos differ for each participant. Throughout experiment runs, we observed common patterns in optimized stimuli across participants, indicating generalizability of stimulus properties. Clarifying the degree to which optimized stimuli generalize to new individuals is important for both clinical applications in neuromodulation and visual neuroscientific understanding. Stimuli that generalize would allow neuromodulatory therapeutics to be delivered in “open-loop”, and provide insights about human visual neural function at the population level. To explore this question, we collected data in a new set of “generalization” participants using stimuli derived from the optimization experiments’ data for Theta 40d and all Alpha dimensionalities.

The stimuli shown to the generalization participants were displayed in open-loop and were a combination of preset stimuli (Preset), the preset stimuli that had the lowest and highest average neural objective across optimization participants (Extreme Presets) and the stimuli that had the lowest and highest neural objective from each individual optimization participant (Extreme Par). The mean neural objective values across generalization participants and repeats and 95% bootstrapped confidence intervals are shown for all stimulus types and experiment versions in Figure 11A. The “Extreme Preset” stimuli were chosen by taking all preset trials across all optimization participants for an experiment version: 6 participants, 25 preset stimuli, and 2 repeats for a total of 300 trials, and averaging the neural responses for each of the 25 preset stimuli across participants and repeats, then choosing the stimuli with highest and lowest average neural objectives. The “Extreme Par” stimuli were chosen independently from each optimization participant’s trials (250 trials) based on the maximum and minimum neural objectives with no averaging. The number of trials going into each decision were comparable, but not equal (300 versus 250) and compare random exploration with within- and across-participant averaging versus participant-specific optimization as methods for generating generalizable stimuli. We find that “Extreme Preset” stimuli do not show significant effect size (Hedges’ *g*) between minimizing (black) and maximizing (red) stimuli for any experiment version, while “Extreme Par” stimuli show significant effect size for all experiment versions: [0.51, 0.68, 1.07, 0.58] for panels left-to-right (*p <* 1e-4, Welch’s t-test, Bonferroni corrected, *n* = 4). This shows that selecting extreme stimuli from a random set with averaging is inefficient for finding generalizable stimuli compared to chosing stimuli optimized for single participants.

**Figure 11:**
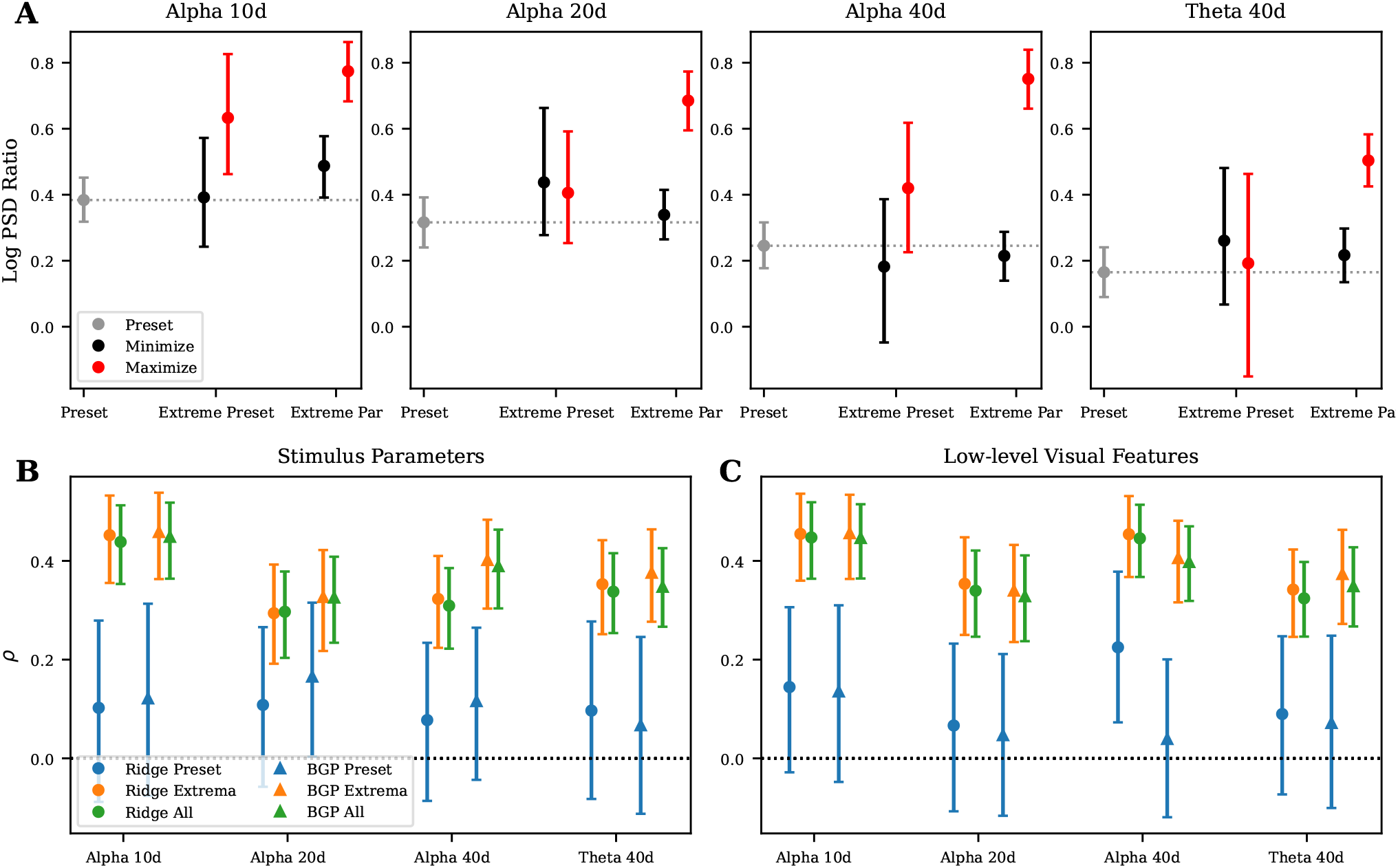
Optimized stimuli generalize to new participants. **A:** Mean and 95% bootstrapped confidence intervals across generalization participants are shown for neural objectives collected from Preset stimuli, preset stimuli with the min/max average neural objective in optimization participants (Extreme Preset), and stimuli with the min/max neural objective from each optimization experiment (Extreme Par) Stimuli from all Alpha dimensions and Theta 40d were collected and are shown across panels. **B:** Mean correlations and 95% bootstrapped confidence intervals on the generalization trials are shown for preset stimuli (blue markers), extreme stimuli (orange markers) and all stimuli (green markers) for crossvalidated Ridge (circles) and Bayesian Gaussian process (BGP) regression (triangles) models trained on the optimization datasets to predict neural objectives from stimulus parameters. **C:** Mean correlations and 95% bootstrapped confidence intervals on the generalization trials are shown for models trained to predict neural objectives from a set of 9 low-level image features computed from the generated texture. Colors and shapes as in **B**.

To further characterize what aspects of neural response functions are generalizable across participants, we trained Ridge and Bayesian Gaussian Process (surrogate model used for closed-loop optimization) encoding models to predict neural objectives from video generation stimulus parameters and low-level visual features. The models were trained on optimization participants’ data and tested on generalization participants’ data. Figure 11B shows the mean linear correlation and 95% bootstrapped confidence intervals between the predicted and ground-truth neural objectives from the generalization participants when the model predictors were the stimulus parameters after softmax and square-root transformation. Both Ridge and Bayesian Gaussian process models struggle to predict the neural objectives for the preset trials, but the Extreme trials are well predicted across models and experiment versions. Similarly accurate predictions are found when the model predictors were a set of 9 low-level visual features (see Image feature extraction for descriptions) extracted from the first frame (texture) of each stimulus video (Fig. 11C). Together, these results suggest that there is some alignment across participants’ neural response functions which allows encoding models to generalize, that linear models perform similarly to non-parametric Bayesian Gaussian processes, and that the modulation in neural objectives are well explained by low-level visual features in additional to the stimulus parameters.

### 2.6 Modulation axes are reliable across participants in lower dimensions

We next sought to understand whether the Ridge encoding model weights (*β*s) had interpretable structure and whether that structure was aligned across optimization participants. To this end, we fit Ridge encoding models to cross- (as in Section 5.5) and within-participant optimization datasets for all optimization experiment versions. Although the linearity assumption of the Ridge model may limit model flexibility to fit the underlying neural response functions, its strong predictive performance comparable to the non-parametric Bayesian Gaussian processes suggests that we can use the linear modulation axis derived from the model to understand neural response alignment across participants and generate a family of textures that evoke low and high neural objectives.

For the Ridge models trained by combining participants in Figure 11B, we can inspect the model weights. Figure 12A shows the model weights (normalized by their standard deviation) for Theta and Alpha (top and bottom) 10d, 20d, and 40d datasets where the predictors are the softmax and squareroot normalized latent texture weights. Across experiment versions, most latent textures weights are negative (black), indicating their contribution to lowering neural objectives. Only a small number of textures have positive weights (red), suggesting that sparse combinations are the main drivers of high log power spectral density ratios. However, there is no clear pattern across neural bands or dimensions. With these weights, we can use the mean of the predictor variables as a starting point, then use the fit regression weights to define a direction in parameter space scanning through stimulus parameter space from the minimum to maximum projections observed in the dataset (see Texture modulation axis for details). This defines a trajectory through the parameter space that is aligned with the low-to-high modulation direction. Figure 12E shows these trajectories in the texture pixel space with low to high predicted neural objectives from left to right. Theta and Alpha 10d show a similar pattern with checkerboards having low predicted neural objectives and the 2nd and 7th texture from Figure 7C having higher predicted neural objectives, respectively.

**Figure 12:**
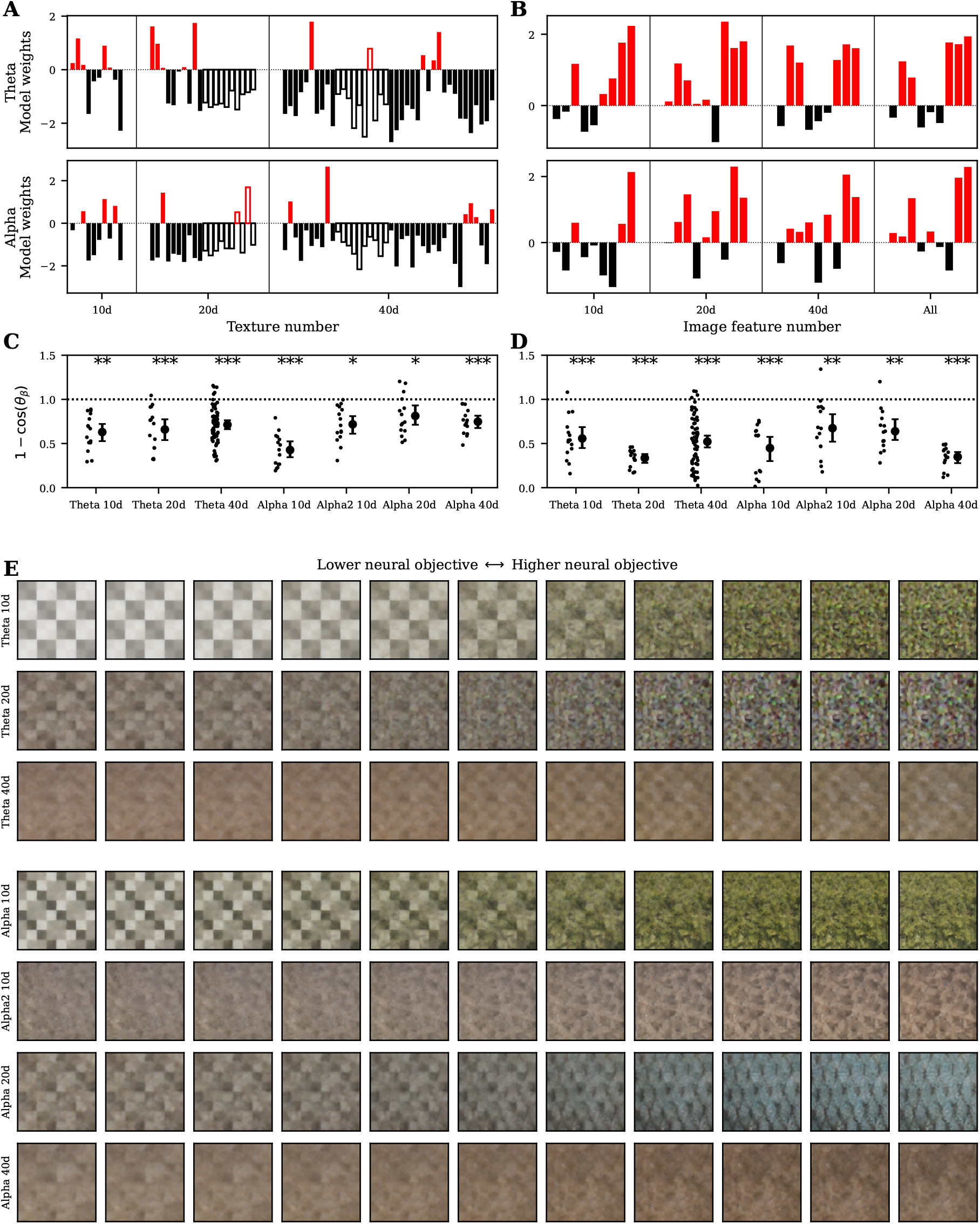
Modulation axes are reliable across participants in lower dimensions. **A:** Ridge regression weights (normalized by their standard deviation) for models trained on 10, 20, and 40d (left to right) and Theta (top) and Alpha (bottom) experiment versions to predict neural objectives from stimulus parameters. The first 10 weights are filled, the second 10 weights are empty bars, and the last 20 weights are filled bars. Red or black bars indicate a positive or negative weights. **B:** Ridge regression weights normalized by their standard deviation for models trained on all Theta (top) and Alpha (bottom) experiment versions for models trained to predict neural objectives from a set of 9 features computed from the generated texture. Colors as in **A. C:** Pairwise cosine distances between participant’s Ridge weights train on stimulus parameters (small dots) and the mean distance and bootstrapped 95% confidence intervals (large dots with errorbars) are shown for all experiment versions. Dashed horizontal line at 1 is the mean of the null distribution assuming randomly distributed model weights. **D:** Pairwise cosine distances between participant’s Ridge weights trained on low-level image features are shown for all experiment versions. Markers, null line, and p-values as in **C. E:** A set of textures spanning the linear encoding subspace of a Ridge model is shown for each experiment version.

**Figure 13:**
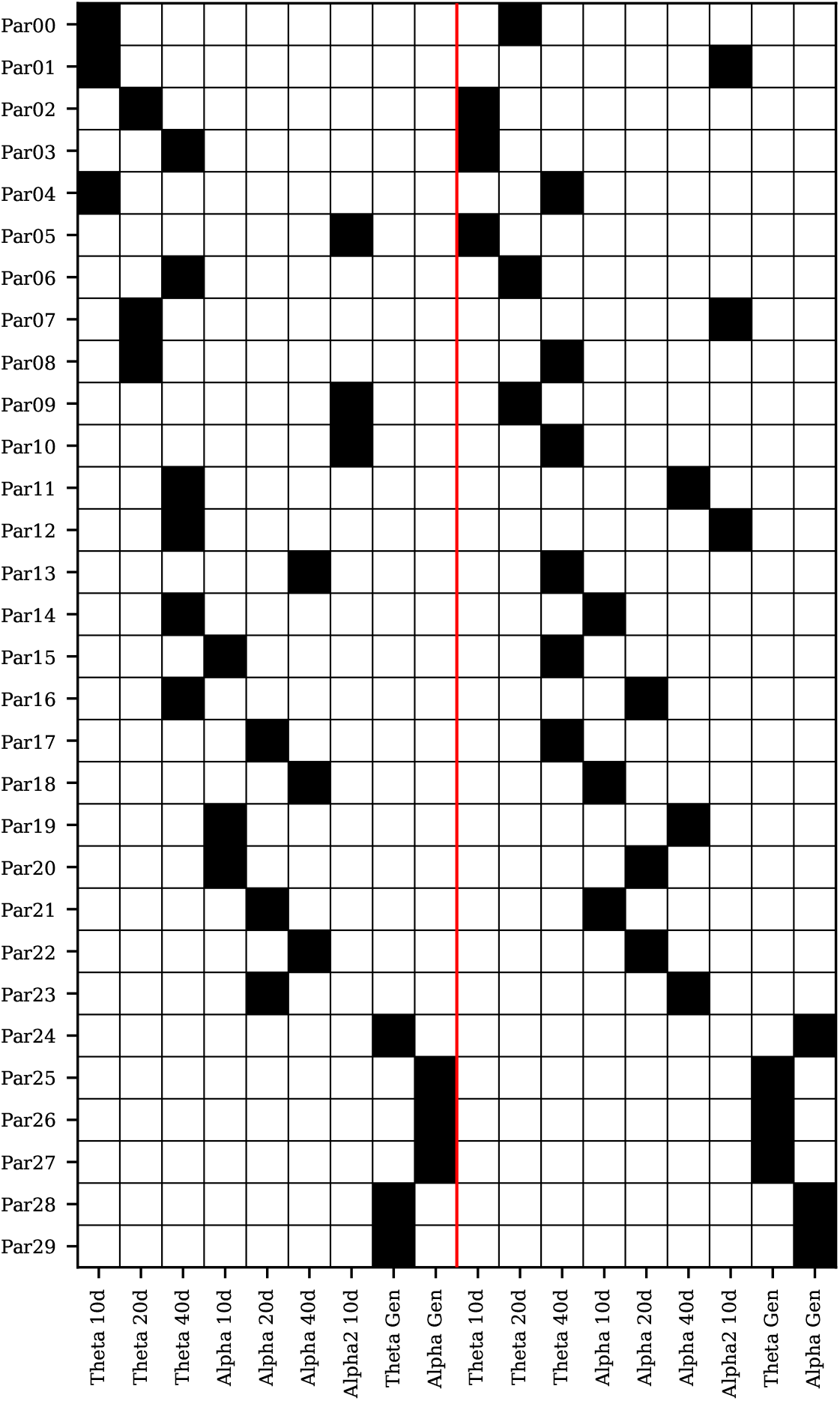
The table shows the mapping between which participants were run with which experiment versions. A black square indicates the participant in the row was run with he experiment version in the column. The experiment versions to the left and right of the vertical red line indicate experiment versions that were run first and second, respectively.

For the Ridge models where the predictors are the low-level visual features (as in Fig. 11C) there is a higher degree of consistency across experiment versions (Fig. 12B). The distance-to-gray and mid and high spatial frequency features are consistently predictive of higher objectives. Interestingly, the low spatially frequency features has relatively large weights and consistently opposite sign across theta and alpha, indicating that this feature is predictive of higher theta power but lower alpha power. Higher dimensional textures are less consistent.

To determine whether there is consistency between participants within experiment versions, we fit Ridge models to each participant’s data within the experiment versions for both stimulus parameters and low-level visual features. Figure 12C,D show the resulting pairwise cosine distances of model weights between optimization experiments for stimulus parameters (C) and low-level image features (D). The scattered points are the individual distances and the mean and 95% bootstrapped confidence intervals are shown to the right of the individual point. For weights trained on stimulus parameters, the lower dimensional experiment versions have smaller pairwise distances. The distances are generally smaller for weights trained on low level image features and do not have a clear pattern across dimensions. Together, this indicates that more consistent linear encoding structure can be estimated from datasets that had lower dimensional stimulus parameter spaces, and consistent structure can be estimated from low-level visual features.

## 3 Methods

### 3.1 Participants

The study was approved by the Biomedical Research Alliance of New York (BRANY) Ethics Committee, and the experiment was performed according to the relevant guidelines and regulations. Written informed consent was obtained from all participants.

A total of 30 participants participated in the study. 15 reported their sex as female, 11 as male, and 4 left the field blank. 27 participants reported their age and the mean, standard deviation, min, and max ages were 51.7, 15.8, 23, and 78, respectively.

### 3.2 Experiment setup

All experiments were performed at the EEG facility of Dandelion Science in Hoboken, New Jersey, United States. EEG data were sampled at 1024 Hz using a BioSemi Active Two amplifier (BioSemi, Amsterdam, Netherlands) from 68 active Ag/AgCl scalp electrodes arranged according to the international standard 10–20 system for electrode placement in a nylon head cap [43]. The Common Mode Sense (CMS) and Driven Right Leg (DRL) electrodes were placed to the left and to the right of POz, respectively. EEG streams were downsampled to 512 Hz with the BioSemi Lab Streaming Layer (LSL) Application.

All stimuli were shown at a distance of 64 cm on an Alienware 27 inch AW2721D monitor with a refresh rate of 120 Hz and resolution of 2, 560 *×* 1, 440. Videos were shown at full monitor height and with a square aspect ratio. Custom Python software based on Psychopy [44] was used to display instructions and stimuli to the participants. In all experiments, participants were instructed to fixate on a red fixation dot at the center of the screen whenever it is present and minimize eye and body movements during trials.

### 3.3 Closed-loop optimization experiments

All optimization experiments were composed of 250 trials with the exception of Alpha2 10d which had 275 trials. Trials were grouped into blocks of 25 with self-paced breaks between blocks. Each trial was composed of 2 seconds of a grey screen with a red fixation dot, 2 seconds of stimulation with a red fixation dot, and 2 second of grey screen without the fixation dot to allow participants to blink and rest their eyes.

The first 25 trials were Preset trials which had a fixed set of stimuli for all participants within an experiment version. Stimulus parameters were shared between preset trials for experiment versions of the same dimensionality, except for the flicker frequency. The remaining 225 trials were an uniform mix of 25 Preset repeats, 50 minimization trials, and 150 maximization trial. The order of trials within each block was randomized per participant. The Alpha2 10d experiment version had an additional 25 trials which used stimuli from the Alpha 10d experiment version to facilitate comparison between the two. After the first 25 preset, closed-loop optimization began with the next required maximization or minimization trial being generated after each new trial was acquired.

Each optimization participant ran two experiment versions with differing dimensionalities and/or neural bands and the order was counter balanced across participants (see Figure A.1 for details). The mean and standard deviation of the experiment run duration for optimization experiments was 29.7 *±* 5.0 minutes.

### 3.4 Open-loop generalization experiments

Each participant for the generalization experiment ran a Theta and Alpha generalization experiment version and the order was counter balanced across participants (see Figure A.1 for details). Trial structure and duration were the same as optimization experiments. The Theta generalization experiment only contained videos from the Theta 40d optimization experiments. The Alpha generalization experiment contained video from Alpha 10d, 20d, and 40d optimization experiments. The generalization experiment versions were composed of an equal number of videos from each source optimization experiment version with the following types

- 25 Preset videos repeated once,
- 2 Extreme Preset videos (1 min, 1 max) repeated 3 times (4 times including the Preset, repeat)
- 2 Extreme Par videos (1 min, 1 max) from 6 optimization participants repeated 4 times, and
- 4 model-based videos repeated 4 times.

The model-based videos were excluded from analysis due to a bug in how the files were saved which was only discovered after the generalization data was collected.

For the theta (40d only) and alpha (all dimensions) generalization experiments, the mean and standard deviations of experiment duration were 12.1 *±* 0.5 and 33.0 *±* 0.9 minutes, respectively.

### 3.5 Video generation

Two-second videos were generated by a combination of a parameterized latent variational autoencoder (VAE) space decoded into image space and sinusoidal dynamics applied in image space. 10, 20, and 40 dimensional latent spaces were created in the latent space of a pretrained VAE taken from a Stable Diffusion image generation model [39]. The Optimizer operated in a [0, 1]^*N*^ parameter space. These parameters were first converted to a unit Gaussian distribution through the inverse cumulative distribution function and clipped to *±* 10. These values were then passed through a softmax function and turned into a point on the standard *n*-simplex (Eq. 13). The softmax-transformed weights were used to create a linear combination of the latent textures(Eq. 14). The combination was passed into the VAE deocder and transformed into a 512 *×* 512 image (Eq. 16,17).

Sinusoidal dynamics at the flicker frequency (6 or 10 Hz) were applied to the images to interpolate them between the image and gray (Eqs. 13).

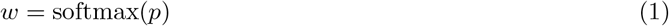

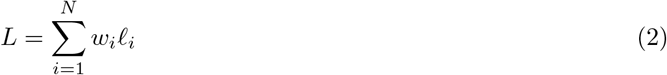

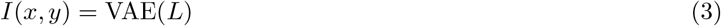

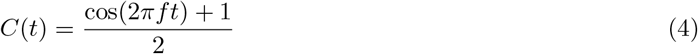

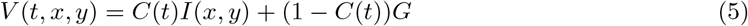

The resulting tensors were clipped to [0, 1], scaled by 255 and converted to integer values before being saved as mp4 files.

Latent textures 1 - 3 were created by sampling independent normal distributions with the VAE latent dimension (4, 64, 64). Latents 1 and 2 were spatially low-pass filtered. Latents 4 and 5 were black and white checkerboard patterns created in image space then passed through the VAE encoder. Latents 6 - 9 were images of natural objects taken by the first author and passed through the VAE encoder. Latent 10 was a white image passed through the VAE encoder.

Latents 11 - 40 were images taken from the Describable Textures Dataset and the Amsterdam Library of Textures [45, 46] cropped square and passed through the VAE encoder. A larger set of images were initially selected and these thirty images were sorted based on least similarity to previously added images computed from average color and distance in VAE latent space. The specific source files are listed in .

### 3.6 EEG preprocessing and neural objective calculation

EEG was windowed to the 2 seconds preceding an LSL trigger which signaled the end of stimulation. All 68 EEG channels were referenced to the common average and 10 occipital and occipital-parietal channels were subselected for neural objective calculation, specifically Oz, O1, O2, PO7, PO3, POz, PO8, PO4, PO9, PO10. The same preprocessing was used for online optimization (MNE-LSL library [47] and offline analysis (MNE-python [48]).

On the 2-second preprocessed signal, a hann window was applied and the power spectral density was calculated for each channel. The bin closest to the flicker frequency was taken as the numerator and the mean of the bins *±* 2 bins away (1 Hz total) was used as the denominator. The log of this ratio was calculated per-channel and the mean across channels was used as the neural objective for optimization for each trial. In this paper, we use “log power spectral density ratio” (log PSD ratio) and “neural objective” interchangeably.

### 3.7 Optimizer and regression models

Bayesian optimization was used with a Bayesian Gaussian process as the surrogate model. Uniform priors were used for the squared-ex√ponential kernel lengthscale, GP variance, and diagonal noise variance, with distributions *U* (1e-3, *d*), *U* (1e-3, 1), and *U* (1e-2, 1), respectively. JAX [49] was used to construct the model and the NUTS sampler from numpyro [50] was used to draw samples from the posterior of the GP parameters.

To select the parameters for the next trial, the posterior samples were used to calculate the posterior predictive distribution, which was calculated per-sample for a set of 8 candidate parameter vectors. The upper confidence bound (UCB) was calculated for each sample from the posterior predictive distribution and the posterior-averaged UCB was calculated. L-BFGS-B [51] was used to optimize the candidate points with respect to the posterior-averaged UCB and the candidate with the highest UCB was selected as the candidate for the next trial. After each new datapoint was added to the optimizer, the neural objectives were re-zscored.

After the first block of preset trials was completed, the NUTS sampler was run for 250 warmup samples. Then, after each additional trial, 250 more steps were taken starting from the last sampler state and the 250 samples were decimated to 50 for use in Bayesian optimization.

### 3.8 Optimization effect size, ICC, confidence intervals, and ANOVA

For both optimization effect size (Hedges’ *g*) and intra-class correlation (ICC), we used the Pingouin Python library [52]. Hedges’ *g* is calculated as

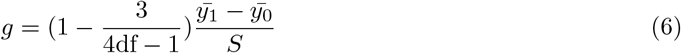

where *S* is the pooled standard deviation and df is the degrees of freedom in the standard deviation. For within-participant ICCs, the 2 repeats of each Preset stimulus are used and ICC(1,1) is reported [42]. For cross-participant ICCs each Preset trial is repeated both within and across participants. In this case, datasets were resampled 1000 times where one of the two within-participant repeats for each preset stimulus was randomly selected for each participant and the median “consistency” ICC(C, 1) was reported, which is invariant to each participant’s mean Preset neural objective. SciPy’s bootstrapping functions were used to calculate bootstrapped confidence intervals with the default bias-corrected and accelerated bootstrap confidence interval method [51]. The ANOVA was performed using the statsmodels Python library’s “ols” and “anova lm” functions.

### 3.9 Regression models

This same Bayesian Gaussian Process surrogate model was used offline as an encoding regression model. All settings were the same except for the number of warm-up samples (1000) and the number of posterior sampled generated (500 which were decimated by a factor of 10).

Ridge and RidgeCV models from scikit-learn [53] were also used for offline encoding regressions since it is straightforward to visualize the axis in predictor space that maximally modulates the predictions. For Ridge models trained on a single participant’s data, the “Leave-One-Out” cross validation strategy was used for the ridge parameter. For models trained on multiple participants’ data, cross-validation was performed leaving one training participant out to select the ridge parameter then the model was re-fit on all training data using the selected ridge parameter. We cross-validate a set of 41 ridge parameters log-spaced between 1 and 10^10^. For all regression models, we report linear correlations rather than the coefficient of determination since cross-validation of the Ridge parameter on neural datasets often leads to large regularization parameters which produces regimes where the model predictions track ground-truth data but at a much smaller absolute variance.

The video generator parameters were softmax transformed and then the square-root was taken (similar to a Hellinger distance [54]) before the parameters were zscored to the training mean and standard deviation for model fitting.

### 3.10 Image feature extraction

Image features extracted from the first frame (texture) of each video were:

1. Luminance: mean of the Lab luminance channel across the image,
2. Contrast: standard deviation of the Lab luminance channel across the image,
3. Distance to gray: average RGB distance to gray across the image,
4. R: average of the R channel across the image,
5. G: average of the G channel across the image,
6. B: average of the B channel across the image,
7. Low-spatial frequency (SF): average log power of the lowest third of log-spaced frequencies (excluding 0),
8. Mid-SF: average log power of the middle third of log-spaced frequencies, and
9. High-SF: average log power of the highest third of log-spaced frequencies (up to the Nyquist frequency of 0.5 cycles/pixel for 512-pixel images).

The low-level image features were zscored to the training mean and standard deviation for model fitting.

### 3.11 Model weight distances

Pairwise model weight distances were calculated using the cosine distance

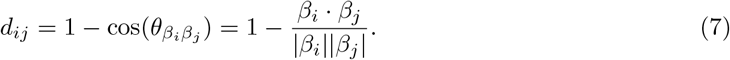

The mean cosine distances in Figure 12C,D are calculated in dimension *d* across *N* participantsleading to 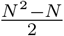 pairwise distances. The null distribution is estimated by drawing *N* random unit vectors in *d* dimensions, computing the pairwise cosine distances, and calculating the mean distance. This is repeated 50,000 times to generate the null distribution of means for calculating the p-values.

### 3.12 Texture modulation axis

For a fitted Ridge regression model, we want to define a trajectory through parameter space that is aligned with the model’s weight coefficient (modulation axis). Stimulus parameter changes parallel to this axis change the predicted neural objective, and stimulus parameter changes orthogonal to this axis leave it unchanged. However, the Ridge models are fit on transformed and zscored stimulus parameters, so we need to first define the trajectory in the linear model and then apply an inverse transform to the parameters. Let *x* and *y* be the transformed data and *x*_*z*_ and *y*_*z*_ be the transformed and zscored data. The fit Ridge model’s predictions look like

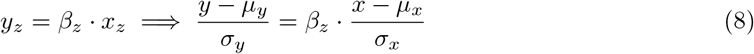

where the *µ* and *σ* variables are the mean and standard deviations of the training features. This can be rearranged to

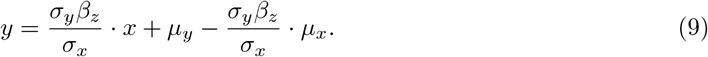

A linear trajectory can be defined by a point and direction. For the point, we use the mean *x* which leads to the mean *y* prediction *y* = *µ*_*y*_:

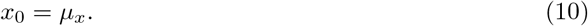

The unit-norm direction is the *β* for the unzscored parameters

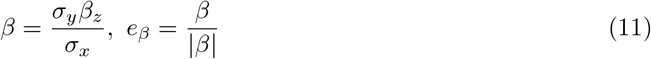

which means the linear trajectory in transformed parameter space is parametrized by

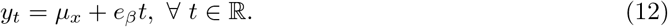

We set the limits of *t* for each experiment version by mean centering all transformed parameters used in the experiment and calculating their projections onto *e*_*β*_. The min and max projections set the limits for *t* and we take 11 linear spaced points along this trajectory for visualization.

To map back into untransformed stimulus parameter space, we square and then take the log of the transformed parameters. These vectors are passed into the video generator and the first frame of each is shown in Figure 12E.

## 4 Introduction

Rhythmic ensemble neural activity (neural oscillations) has been found to robustly associate with perceptual processing, behaviors and mental states. In visual processing, the alpha (8-14 Hz) oscillation has been shown to phase-lock to saccades during visual scene encoding [1], and visual attention in both human and macaque monkeys is modulated by theta (4-7 Hz) [2, 3]. In speech processing, delta (1-4 Hz), alpha, and theta neural oscillations, measurable with EEG, track both acoustic and linguistic features in the speech signal [4]. Beta oscillations (13-30 Hz) and their coupling are shown to be involved in motor preparation [5, 6] and maintenance-of-state [7]. Meanwhile, abnormalities in neural oscillations are shown to correlate with dysfunctions in sensory, motor, and cognitive processes (see review by [8]), leading to a growing body of research on invasive and noninvasive interventions via the modulation of oscillatory activities. For example, neuromodulation of gamma-band activity using visual stimulation has been indicated as a potential tool for slowing down cognitive decline in older subjects and patients with Alzheimer’s disease [9, 10, 11]. Non-invasive electrical stimulation has been suggested as a method for partially restoring vision with activity in the alpha band as a specific neuromodulatory target [12]. However, existing techniques for modulating oscillations are highly constrained in their parameter space, for example, by only changing the amplitude (intensity) of the stimulation, thus cannot be easily optimized or adapted to individuals, resulting in high cross-subject differences in their outcomes [13, 14, 15, 16].

One promising approach for modulating neural oscillations in individual subjects is closed-loop neural optimization. To achieve this, a predefined “neural objective” in response to the input stimulus at trial *t* is fed back to a stimulus synthesizer to alter the parameter for the input at trial *t* + 1, in order to alter the neural feature in a designated direction, e.g., maximizing the amplitude of alpha-band activity at a certain electrode. Such a realtime closed-loop approach has been successfully utilized in visual [17, 18, 19] and motor [20, 21] systems in animal models. However, application in human subjects remain limited largely due to the poor signal-to-noise ratio in noninvasive measurements especially from relatively low-cost devices such as electroencephalography (EEG) [22, 23].

Some existing studies of closed-loop EEG optimization [24, 25] require the participant’s active but implicit effort, making the optimization procedure long and results less interpretable and generalizable [26]. Moreover, previous stimulus optimization studies often operate in a very low dimensional space where it is possible to test a regular grid of parameters, which is not feasible in even 10s of dimensions. For example, combinations of color or luminance and frequency within the gamma band for entrainment [27] or shape size and contrast for modulation of endogenous theta[28]. Even when generative models were used for stimulus generation, the stimuli produced vary along simple dimensions of grayscale luminance and basic shapes [23].

In this study, we establish the feasibility of a closed-loop visual stimulus optimization pipeline using widely available EEG hardware and passive video viewing, readily exploitable for both fundamental and clinical research. We implemented and validated a closed-loop system that optimizes human rhythmic EEG response to visual textures parametrized in the latent space of a deep generative image model. The generative image model we use can generate high dimensional (up to ∼ 16, 000) naturalistic color images and textures, although here we parametrically constrain the search space to a subspace of the full dimensionality to understand the dependence of optimization effect on dimensionality. Our neural objective is based on steady-state visual evoked potentials (SSVEP) features, a common technique used in visual neuroscience [29, 30, 31, 32] and braincomputer interface research [33, 34, 35] to boost the signal-to-noise ratio of EEG-based features. We show that stimulus parameters that maximize and minimize our neural objective can be identified within a single 30-minute session in a 10-dimensional search space. We then show that the same system is also able to optimize in 20- and 40-dimensional spaces, although the optimization effect size, which we quantify with Hedges’ *g* (an unbiased version of the more familiar Cohen’s *d* [36, 37]), is smaller for larger dimensionalities. At the individual participant level, participant-specific factors explain more of the observed variability in optimization effect size than dimensionality or neural band. Next, we demonstrate that the stimulus latent parameters and extracted low-level visual features can both be used to train encoding models to predict neural objectives in new participants. The distribution of power in spatial frequencies is the most consistent predictor of neural objectives with low spatial frequencies having opposite effects on theta and alpha bands. Finally, we visualize trajectories in stimulus space that maximally modulate the neural objective for all dimensionalities and both neural bands. Together, these results show that closed-loop optimization in humans using EEG is a viable approach for producing stimuli optimized to drive neural targets of neuroscientific and neuromodulatory interest.

## 5 Results

A total of 30 subjects participated in two types of electroencephalography (EEG) experiments without overlapping: *optimization* (*n* = 24) and *generalization* (*n* = 6). In both types of experiments, we defined the “neural objective” as the log ratio of the spectral power density (PSD) of the EEG signal at theta (6 Hz) or alpha (10 Hz) flicker frequency to the adjacent frequency bins averaged across 10 occipital and occipital-parietal electrodes in the same 2 second window as the stimulus presentation. In *optimization* experiments, subjects were instructed to passively watch short flickering videos while their real-time EEG response was used to generate new videos for optimizing, that is, maximizing or minimizing, their neural objectives using our closed-loop optimization system. Videos generated in optimization experiments were pooled and filtered for the *generalization* experiments, where participants were similarly instructed to passively watch short video clips, but the videos were predefined and did not adapt to their EEG response.

In *optimization* experiments, three different dimensionalities of stimulus parameter (texture) space were tested at each flicker frequency: 10, 20, and 40. In *generalization* experiments, all three dimensionalities were tested at 10 Hz whereas only 40-dimensional stimuli were tested at 6 Hz. We refer to each combination of experiment type, flicker frequency, and stimulus parameter dimension, as an experiment *version*. For *optimization* experiments, each participant was tested for two separate experiment runs, each consisting of a single version without repetition. All experiment versions were run in 6 participant except for Theta 40d which was run in 12 participants although only the first 6 participants were used for generalization. For *generalization* experiments, all versions of the same flicker frequency were aggregated in the same experiment run. Each experiment version lasted approximately 30 minutes or less.

### 5.1 Closed-loop stimulus optimization setup

Each optimization experiment was initiated with 25 “preset” trials which were used to warm-start optimization and common to all participants within an experiment version. The remaining 225 trials were a mix of 150 “maximization” trials, 50 “minimization” trials, and 25 trials repeating the preset trials. The 225 trials were evenly split into 25-trial blocks and the within-block order was pseudorandomized per-participant, maintaining a roughly constant fraction of each type in every 25 trial block (Fig. 7**A**). Note that the fraction denotes the average number of trials in each block, for example a total of 25 preset trials are uniformly distributed among the 26th to 250th trials, but the exact number of preset trials varies between blocks since 25 does not divide evenly into 9. Trials consist of 2 seconds of fixation, 2 seconds of stimulation, and 2 seconds of rest (Fig. 7**B**). The optimization space consisted of a softmax-weighted linear combination of 10, 20, or 40 latent textures. Fig. 7**C** shows these latent textures projected onto image space with the top row, two rows, and all rows being used for 10, 20, and 40d optimization, respectively. The same 10, 20 and 40d textures were collected for both flicker frequencies. In addition, a second Alpha 10d set of experiments were collected, referred to as “Alpha2 10d” with a different set of 10 textures outlined in red. Extended data Figure 13 shows which participants ran which experiment versions. See Section 6 for more details on the experiment design.

Figure 7D shows a diagram of the closed-loop experimental structure. Starting at the bottom and proceeding in a clockwise direction: first, stimulus parameters, *p*, are converted to weights, *w* by a softmax operation. These weights are used to form a linear combination of the latent textures in Figure 7**C**. The latent texture is passed through the decoder of a variational autoencoder (VAE) taken from a latent diffusion model [38, 39] to create an image, and the image pixels are modulated sinusoidally, at the given flicker frequency, between the image and a uniform gray background to produce a video. Second, the video is displayed to the participant while the raw EEG signal is streamed through Lab Stream Layer (LSL) [40]. Third, the raw EEG undergoes minimal online preprocessing and the log power spectral density ratio neural objective is calculated for the occipital and occipital-parietal channels shown in red in the topoplot (see Section 6.6 for details). Finally, the paired stimulus parameters and objective are added to the trial history and a Bayesian optimizer is used to select the next set of parameters to test. This process is repeated throughout the duration of the experiment.

### 5.2 Visual stimulation robustly evokes SSVEP for Theta and Alpha 10d

Periodic flickering of visual stimulus features is a commonly used technique to evoke steady-state visual evoked potentials (SSVEP) [29], which manifest as sharp peaks in neural power spectral densities at the flicker frequency and its harmonics. Low-level visual stimulus features have been found to modulate the amplitude and phase of SSVEP [30]. Here, we first sought to show that our stimulus optimization space could be used to controllably and differentially evoke SSVEP. For minimization and maximization trials, the optimized videos were able to differentially evoke SSVEP amplitude in occipital electrode. Figure 8A and B show the median SSVEP for minimization trials and maximization trials, respectively, for the participant that had the highest optimization effect in the Theta 10d experiment (Par04). The maximization trials show entrainment to the 6 Hz flicker stimulation starting just after stimulus onset (*t* = 0) and subsiding one cycle after stimulus offset (*t* = 2).

We next looked to characterize the spectral and spatial characteristics of the SSVEP signal. We found that the SSVEP is spectrally localized to the flicker frequency and its harmonics with small amounts of evoked power leaking into neighboring frequency bins. Figure 8C plots the log of the power spectral density ratio between the flicker frequency and the average of the 2 frequency bins *±* 2 bins away around the flicker frequency, averaged across trials and EEG electrodes used for computing the neural objective. The stimulation period was 2 seconds, therefore the frequency resolution is limited to 0.5 Hz. The maximization trials show a significantly larger mean log power spectral density ratio compared to the minimization trials at the flicker frequency (difference in mean is 0.29, Welch’s t-test, *p <* 1e-17, df = 93.0). Figure 8D,E show the mean per-electrode log power spectral density ratio taken over minimization and maximization trials, respectively. Spatially, high log power spectral density ratio values are clustered over occipital electrodes, with smaller overall amplitudes in minimization trials compared to maximization trials. Across participants, the mean and standard deviation of the neural objective for maximization trials mainly peaked around occipital electrodes (Fig. 8F,G).

Qualitatively, the SSVEP looks similar for the participant (Par21) that had the highest optimization effect size for Alpha 10d (Fig. 8H-N). The maximization trials show a significantly larger mean log power spectral density ratio compared to the minimization trials at the flicker frequency (difference in mean is 0.37, Welch’s t-test, *p <* 1e-26, df = 82.4). The log power spectral density ratio is strongest over occipital electrodes, although this participant shows a higher degree of lateral asymmetry than the Theta 10d participant (Fig. 8K-L). The cross-participant mean and standard deviations of the means both show structure in occipital electrodes (Fig. 8M,N). Together, these results demonstrate that the flicker-induced SSVEP is spectrally and spatially localized for both Theta and Alpha 10d, and that SSVEP amplitude varies across different optimization trial types in a controlled fashion.

### 5.3 SSVEP is reliably optimized for Theta and Alpha 10d

For optimization to be considered successful, it must be able to drive the distribution of evoked neural objectives apart between minimization versus maximization trials: the farther apart the distributions, the more effective the optimization is. Here, we quantify the optimization effect size by Hedges’ *g*, which is derived from Cohen’s *d* with a bias adjustment for small sample sizes [37].

Figure 9A,B show neural objectives recorded in each trial for the participants with highest optimization effect sizes for Theta and Alpha 10d, participants Par04 and Par21 respectively. Visually, objectives in maximization and minimization separate after about 75 trials in both participants. To quantify the separation across all participants, we aggregate and histogram all trials. Figure 9C,D show histograms of the neural objectives for preset (gray), minimization (black), and maximization trials (red) for trials taken from all participants for Theta and Alpha, respectively. The mean neural objective in the maximization trials is significantly higher than those in the minimization trials (difference in means is 0.49 for Theta 10d *p <* 1e-52, df = 619.6; difference in means is 0.43 for Alpha 10d *p <* 1e-49, df = 721.7). For both flicker frequencies the mean preset neural objective falls between the minimization and maximization trials. This shows that optimized stimuli are able to modulate the distribution of neural objectives between minimization and maximization trials for Theta and Alpha 10d.

In general, Bayesian optimization methods trade-off so-called exploration and exploitation at a trial-by-trial basis, testing places where the surrogate model (see Section 6.7 for more detail) has high uncertainty as quantified by the predicted standard deviation (“exploration”) and places where the surrogate model predicts good objective values as quantified by the predicted mean (“exploitation”). In particular, we use the upper confidence bound acquisition function to adjudicate between candidate parameters, which selects points that have high (or low for minimization) predicted means and large standard deviations. This tradeoff and the single-trial variability of the neural recordings mean that the per-trial optimization effect size is not expected to strictly monotonically increase. To quantify how optimization unfolds across trials, we calculate the cumulative optimization effect size at each trial number, taking all minimization and maximization trials up to that point into account (Fig. 9E,F). Each colored line represents an individual participant, and the black line combines trials from all participants. For Theta 10d, the effect size tends to increase across trials and starts to plateau around 225 trials for some participants. For Alpha 10d, 3 of the participants’ effect sizes generally increase over trials (Par18, Par15, Par21), 2 participants’ peak around the 100th trials (Par14, Par20). However, 1 participant’s effect size hovers around 0 (Par19), indicating unsuccessful optimization. Note the effect sizes estimated with small numbers of trials (near the left extreme of the plot) are likely unreliable since they are estimated from a small number of trials. By the final trial, all Theta 10d participants have significant optimization effect sizes greater than 0.5 (*p <* 1e-6, Bonferroni-corrected Welch’s t-test, *n* = 6) and 5 of 6 Alpha 10d participants have effect sizes greater then 0.5 (*p <* 1e-6, Bonferroni-corrected Welch’s t-test, *n* = 6) and 1 with an effect size near zero. Together, the aggregated and trial-wise optimization effect sizes both indicate that Theta and Alpha SSVEP can indeed be reliably optimized in a 10d latent texture space.

### 5.4 Optimization effect size depends on experiment version and individual participant

Higher-dimensional stimulus parameter search spaces would allow more varied visual statistics to be searched over. Successful optimization in higher dimensional stimulus parameter spaces would enable a more detailed understanding of visual function and stronger and more targeted neuromodulatory effects. However, blackbox optimization will only scale so far before the exponential growth of the search space as a function of dimensionality makes optimization exponentially more difficult: the “curse of dimensionality” [41]. It is not currently known how closed-loop, blackbox optimization will scale as a function of parameter space dimensionality in humans with EEG-based neural objectives.

In order to determine how optimization scales with stimulus parameter dimensionality, we ran optimization experiments with a fixed trial budget in latent texture spaces of 10, 20, and 40 dimensions. In addition, we ran a second set of Alpha 10d experiments with a set of 10 non-overlapping latent textures (Alpha2 10d). Figure10A summarizes the distributions of neural objectives recorded in all versions of optimization experiments. Gray, black, and red violin plots show the distribution of preset, minimization, maximization trials, respectively and are grouped horizontally by experiment version. For the Alpha2 10d experiment version, 25 preset trials from the main Alpha 10d experiment were included in addition to the Alpha2 10d preset trials to aid in comparing optimization at fixed dimension across latent textures. The corresponding neural objectives are summarized separately in the left and right gray violin plots in the Alpha2 10d group, respectively. For all experiment versions, maximization trials have a significantly higher mean than minimization or preset trials (*p <* 1e-6, Welch’s t-test, Bonferroni-corrected with *n* = 14).

Conceptually, there are two main effects that determine the success of blackbox optimization as a function of parameter dimensionality. Figure 10B illustrates three hypothetical sets of scaling curves. As the search space grows, the optimal stimulus in the search space can only get closer to the globally optimal stimulus (or reach a plateau). This means that the potential for optimization effect size (*P* (*d*)) should monotonically increase as a function of dimensionality (Fig. 10B, top panel). The competing effect is that the amount of information gained per trial (*I*(*d*)) about the stimulus-response function goes down as the search space grows (Fig. 10B, middle panel). This is the curse of dimensionality. Together, these two effects mean that optimization effect size (proportional to the product *P* (*d*) *× I*(*d*), Fig. 10,B bottom panel) should increase to a peak at some dimensionality that depends on the specific shapes of these two effects, then decrease as the curse of dimensionality overtakes the benefit of increasing the stimulus parameter space dimensionality. In the red example, the growing stimulus space quickly saturates (*P* (*d*)) and the information gained per trial (*I*(*d*)) drops quickly, together leading to a sharp increase then decrease in optimizer effectiveness around 5d. In the brown example, *P* (*d*) does not saturate over the whole range and *I*(*d*) has a much slower drop, leading to an optimization effectiveness curve that increases up to 40d before starting to decrease. The purple example falls between these two. The details of each experimental design and participant will potentially have its own set of scaling curves which could depend on factors including how the search space manifold is constructed at each dimensionality, individual differences in structural or functional organization of participants’ brains, and the quality of the neural recording.

In order to determine how optimization scales with parameter space dimensionality in the paradigm presented here, we ran experiment versions with 10, 20, and 40 dimensional stimulus parameter spaces. For all experiment versions except for Alpha2 10d, the smaller stimulus parameter spaces are subspaces of the larger parameter spaces and are the same for theta and alpha bands (Figure 7). Figure 10C shows optimization effect sizes and 95% bootstrapped confidence intervals for all experiment versions, as quantified by Hedges’ *g* computed by combining all minimization and maximization trials across participants. Note that the left and right Alpha 10d markers are for Alpha 10d and Alpha2 10d, respectively. We find that all experiment versions have meaningful and statistically significant optimization effect sizes (*p <* 2e − 5, Bonferroni corrected t-tests, *n* = 7). Effect sizes range from 0.6 to 1.0 for all experiment versions except for Alpha 40d where it is 0.32. For the main set of Theta and Alpha experiment versions (excluding Alpha2 10d) the effect size decreases as a function of dimensionality: {10d: 1.00, 20d: 0.85, 40d: 0.70} and 10d:{ 0.99, 20d: 0.73, 40d: 0.32 }, respectively. This indicates that the main set of experiment versions are in the dimensionality regime where optimization can still be reliably successful, but increased dimensionality beyond 10d makes closed-loop optimization less effective.

For the main set of experiment versions, we had to make an arbitrary choice about which textures to start with in 10d and how to grow our search parameter space. This leaves the question of whether a different choice of how to structure the search parameter space could have led to a different scaling across dimensionalities. To partially answer this question, we created the Alpha2 10d experiment version, which uses 10 textures that are only in the Alpha 20d and 40d experiment versions. Alpha2 10d has an optimization effect size of 0.60 (Fig. 10C, Alpha 10d marker on the right), which is lower than both the Alpha 10d and 20d experiment versions. One possible explanation for this is that the latent textures chosen for the Alpha 10d experiment are closer to the optimal stimulus compared to those chosen for the Alpha2 10d experiment, which means that for the same experiment paradigm, specific choices such as how to grow the parameter search space can impact optimization effect size scaling by altering *P* (*d*) (Fig. 10B) in this case.

Unlike open-loop experiments where all participants can be shown the same set of stimuli, closedloop experiments adapt to individual participants. This can potentially lead to more varied results per-participant since optimization outcomes are sensitive to both cross-participant variability and and the cross-trial dynamics of Bayesian optimization. When broken-out by participant, we find that there is substantial cross-participant variability in optimization effect size. Figure 10D shows the per-participant optimization effect size across all experiment versions. Marker shape and horizontal-axis order is the same as in Figure 10C. Here, marker color and fill indicate Benjamini-Hochberg false discovery rate corrected p-values with filled black points indicating *p <* 0.01, filled gray points for 0.05 *> p* ≥0.01, and open gray points for *p* ≥0.05. The spread in per-participant optimization effect sizes is large compared to the confidence intervals estimated for the experiment version-level optimization effect sizes, indicating that participant-specific factors may explain some of the variability in each experiment version.

We next sought to explain this variability in optimization effect sizes across participants and experiment versions. For individual participants within an experiment version, the preset trials are a shared set of trials which can be used to estimate overall SSVEP susceptibility through the preset objective mean and how the cross-stimulus variance is partitioned between noise and structure through an intra-class correlation (ICC)[42]. Across participants, there is substantial variability in the distribution of neural objectives on the preset trials. Figure 10E shows the mean (horizontal axis) and standard deviation (vertical axis) for individual participants for each experiment version (small unfilled markers) and the means of these quantities across participants for each experiment version (large solid markers). The correlation across all participants and experiment versions (*ρ*_*p*_ = 0.49, *p* = 1.0e-3, *n* = 42) and for averages within experiment versions (*ρ*_*v*_ = 0.91, *p* = 1.3e-2, *n* = 6) are significant. Across participants and experiment versions, the preset trial neural objective mean is correlated with the optimization effect size (Fig. 10F, *ρ*_*p*_ = 0.40, *p* = 8.5e-3, *n* = 42) and across experiment versions there is a medium but statistically underpowered correlation(*ρ*_*v*_ = 0.49, *p* = 3.3e-1, *n* = 6). Figure 10G shows the optimization effect size scattered against the reliability of the preset objectives as quantified by ICC(1,1) for individual participants and ICC(C,1) for each experiment version. The correlation across all participants and experiment versions (*ρ*_*p*_ = 0.32, *p* = 4.1e-2, *n* = 42) is significant, and the correlation across experiment versions is large but statistically underpowered (*ρ*_*v*_ = 0.76, *p* = 7.9e-2, *n* = 6). In order to disentangle the impact of the participant and experiment version factors, we ran a Type 2 ANOVA to determine the effect that preset mean, ICC, log-parameter dimensionality, and binary categorical neural band have on optimization effect size. We found that there was an overall statistically significant effect (*F* = 3.83, *p* = 0.011, see A.2 for more details). While the variances explained by individual factors are generally not significant, we note that participant-specific factors have the largest sum-of-squares followed by the log-parameter dimensionality. The neural band has a negligible sum-of-squares. Together, this shows that participant-specific factors explain the majority of optimization effect size variability at the participant level.

### 5.5 Optimized stimuli generalize to new participants

Although the optimization effect size shows a high degree of variability across individual participants, it is not clear to what degree neural response functions are aligned across participants since the generated videos differ for each participant. Throughout experiment runs, we observed common patterns in optimized stimuli across participants, indicating generalizability of stimulus properties. Clarifying the degree to which optimized stimuli generalize to new individuals is important for both clinical applications in neuromodulation and visual neuroscientific understanding. Stimuli that generalize would allow neuromodulatory therapeutics to be delivered in “open-loop”, and provide insights about human visual neural function at the population level. To explore this question, we collected data in a new set of “generalization” participants using stimuli derived from the optimization experiments’ data for Theta 40d and all Alpha dimensionalities.

The stimuli shown to the generalization participants were displayed in open-loop and were a combination of preset stimuli (Preset), the preset stimuli that had the lowest and highest average neural objective across optimization participants (Extreme Presets) and the stimuli that had the lowest and highest neural objective from each individual optimization participant (Extreme Par). The mean neural objective values across generalization participants and repeats and 95% bootstrapped confidence intervals are shown for all stimulus types and experiment versions in Figure 11A. The “Extreme Preset” stimuli were chosen by taking all preset trials across all optimization participants for an experiment version: 6 participants, 25 preset stimuli, and 2 repeats for a total of 300 trials, and averaging the neural responses for each of the 25 preset stimuli across participants and repeats, then choosing the stimuli with highest and lowest average neural objectives. The “Extreme Par” stimuli were chosen independently from each optimization participant’s trials (250 trials) based on the maximum and minimum neural objectives with no averaging. The number of trials going into each decision were comparable, but not equal (300 versus 250) and compare random exploration with within- and across-participant averaging versus participant-specific optimization as methods for generating generalizable stimuli. We find that “Extreme Preset” stimuli do not show significant effect size (Hedges’ *g*) between minimizing (black) and maximizing (red) stimuli for any experiment version, while “Extreme Par” stimuli show significant effect size for all experiment versions: [0.51, 0.68, 1.07, 0.58] for panels left-to-right (*p <* 1e-4, Welch’s t-test, Bonferroni corrected, *n* = 4). This shows that selecting extreme stimuli from a random set with averaging is inefficient for finding generalizable stimuli compared to chosing stimuli optimized for single participants.

To further characterize what aspects of neural response functions are generalizable across participants, we trained Ridge and Bayesian Gaussian Process (surrogate model used for closed-loop optimization) encoding models to predict neural objectives from video generation stimulus parameters and low-level visual features. The models were trained on optimization participants’ data and tested on generalization participants’ data. Figure 11B shows the mean linear correlation and 95% bootstrapped confidence intervals between the predicted and ground-truth neural objectives from the generalization participants when the model predictors were the stimulus parameters after softmax and square-root transformation. Both Ridge and Bayesian Gaussian process models struggle to predict the neural objectives for the preset trials, but the Extreme trials are well predicted across models and experiment versions. Similarly accurate predictions are found when the model predictors were a set of 9 low-level visual features (see Section 6.10 for descriptions) extracted from the first frame (texture) of each stimulus video (Fig. 11C). Together, these results suggest that there is some alignment across participants’ neural response functions which allows encoding models to generalize, that linear models perform similarly to non-parametric Bayesian Gaussian processes, and that the modulation in neural objectives are well explained by low-level visual features in additional to the stimulus parameters.

### 5.6 Modulation axes are reliable across participants in lower dimensions

We next sought to understand whether the Ridge encoding model weights (*β*s) had interpretable structure and whether that structure was aligned across optimization participants. To this end, we fit Ridge encoding models to cross(as in Section 5.5) and within-participant optimization datasets for all optimization experiment versions. Although the linearity assumption of the Ridge model may limit model flexibility to fit the underlying neural response functions, its strong predictive performance comparable to the non-parametric Bayesian Gaussian processes suggests that we can use the linear modulation axis derived from the model to understand neural response alignment across participants and generate a family of textures that evoke low and high neural objectives.

For the Ridge models trained by combining participants in Figure 11B, we can inspect the model weights. Figure 12A shows the model weights (normalized by their standard deviation) for Theta and Alpha (top and bottom) 10d, 20d, and 40d datasets where the predictors are the softmax and square-root normalized latent texture weights. Across experiment versions, most latent textures weights are negative (black), indicating their contribution to lowering neural objectives. Only a small number of textures have positive weights (red), suggesting that sparse combinations are the main drivers of high log power spectral density ratios. However, there is no clear pattern across neural bands or dimensions. With these weights, we can use the mean of the predictor variables as a starting point, then use the fit regression weights to define a direction in parameter space scanning through stimulus parameter space from the minimum to maximum projections observed in the dataset (see Section 6.12 for details). This defines a trajectory through the parameter space that is aligned with the low-to-high modulation direction. Figure 12E shows these trajectories in the texture pixel space with low to high predicted neural objectives from left to right. Theta and Alpha 10d show a similar pattern with checkerboards having low predicted neural objectives and the 2nd and 7th texture from Figure 7C having higher predicted neural objectives, respectively.

For the Ridge models where the predictors are the low-level visual features (as in Fig. 11C) there is a higher degree of consistency across experiment versions (Fig. 12B). The distance-to-gray and mid and high spatial frequency features are consistently predictive of higher objectives. Interestingly, the low spatially frequency features has relatively large weights and consistently opposite sign across theta and alpha, indicating that this feature is predictive of higher theta power but lower alpha power. Higher dimensional textures are less consistent.

To determine whether there is consistency between participants within experiment versions, we fit Ridge models to each participant’s data within the experiment versions for both stimulus parameters and low-level visual features. Figure 12C,D show the resulting pairwise cosine distances of model weights between optimization experiments for stimulus parameters (C) and low-level image features (D). The scattered points are the individual distances and the mean and 95% bootstrapped confidence intervals are shown to the right of the individual point. For weights trained on stimulus parameters, the lower dimensional experiment versions have smaller pairwise distances. The distances are generally smaller for weights trained on low level image features and do not have a clear pattern across dimensions. Together, this indicates that more consistent linear encoding structure can be estimated from datasets that had lower dimensional stimulus parameter spaces, and consistent structure can be estimated from low-level visual features.

## 6 Methods

### 6.1 Participants

The study was approved by the Biomedical Research Alliance of New York (BRANY) Ethics Committee, and the experiment was performed according to the relevant guidelines and regulations. Written informed consent was obtained from all participants.

A total of 30 participants participated in the study. 15 reported their sex as female, 11 as male, and 4 left the field blank. 27 participants reported their age and the mean, standard deviation, min, and max ages were 51.7, 15.8, 23, and 78, respectively.

### 6.2 Experiment setup

All experiments were performed at the EEG facility of Dandelion Science in Hoboken, New Jersey, United States. EEG data were sampled at 1024 Hz using a BioSemi Active Two amplifier (BioSemi, Amsterdam, Netherlands) from 68 active Ag/AgCl scalp electrodes arranged according to the international standard 10–20 system for electrode placement in a nylon head cap [43]. The Common Mode Sense (CMS) and Driven Right Leg (DRL) electrodes were placed to the left and to the right of POz, respectively. EEG streams were downsampled to 512 Hz with the BioSemi Lab Streaming Layer (LSL) Application.

All stimuli were shown at a distance of 64 cm on an Alienware 27 inch AW2721D monitor with a refresh rate of 120 Hz and resolution of 2, 560 *×* 1, 440. Videos were shown at full monitor height and with a square aspect ratio. Custom Python software based on Psychopy [44] was used to display instructions and stimuli to the participants. In all experiments, participants were instructed to fixate on a red fixation dot at the center of the screen whenever it is present and minimize eye and body movements during trials.

### 6.3 Closed-loop optimization experiments

All optimization experiments were composed of 250 trials with the exception of Alpha2 10d which had 275 trials. Trials were grouped into blocks of 25 with self-paced breaks between blocks. Each trial was composed of 2 seconds of a grey screen with a red fixation dot, 2 seconds of stimulation with a red fixation dot, and 2 second of grey screen without the fixation dot to allow participants to blink and rest their eyes.

The first 25 trials were Preset trials which had a fixed set of stimuli for all participants within an experiment version. Stimulus parameters were shared between preset trials for experiment versions of the same dimensionality, except for the flicker frequency. The remaining 225 trials were a uniform mix of 25 Preset repeats, 50 minimization trials, and 150 maximization trials. The order of trials within each block was randomized per participant. The Alpha2 10d experiment version had an additional 25 trials which used stimuli from the Alpha 10d experiment version to facilitate comparison between the two. After the first 25 preset, closed-loop optimization began with the next required maximization or minimization trial being generated after each new trial was acquired.

Each optimization participant ran two experiment versions with differing dimensionalities and/or neural bands and the order was counter balanced across participants (see Figure A.1 for details). The mean and standard deviation of the experiment run duration for optimization experiments was 29.7 *±* 5.0 minutes.

### 6.4 Open-loop generalization experiments

Each participant for the generalization experiment ran a Theta and Alpha generalization experiment version and the order was counter balanced across participants (see Figure A.1 for details). Trial structure and duration were the same as optimization experiments. The Theta generalization experiment only contained videos from the Theta 40d optimization experiments. The Alpha generalization experiment contained video from Alpha 10d, 20d, and 40d optimization experiments. The generalization experiment versions were composed of an equal number of videos from each source optimization experiment version with the following types

- 25 Preset videos repeated once,
- 2 Extreme Preset videos (1 min, 1 max) repeated 3 times (4 times including the Preset, repeat)
- 2 Extreme Par videos (1 min, 1 max) from 6 optimization participants repeated 4 times, and
- 4 model-based videos repeated 4 times.

The model-based videos were excluded from analysis due to a bug in how the video files were saved which was only discovered after the generalization data was collected.

For the theta (40d only) and alpha (all dimensions) generalization experiments, the mean and standard deviations of experiment duration were 12.1 *±* 0.5 and 33.0 *±* 0.9 minutes, respectively.

### 6.5 Video generation

Two-second videos were generated by a combination of a parameterized latent variational autoencoder (VAE) space decoded into image space and sinusoidal dynamics applied in image space. 10, 20, and 40 dimensional latent spaces were created in the latent space of a pretrained VAE taken from a Stable Diffusion image generation model [39]. The Optimizer operated in a [0, 1]^*N*^ parameter space. These parameters were first converted to a unit Gaussian distribution through the inverse cumulative distribution function and clipped to *±* 10. These values were then passed through a softmax function and turned into a point on the standard *n*-simplex (Eq. 13). The softmax-transformed weights were used to create a linear combination of the latent textures(Eq. 14). The combination was passed into the VAE decoder and transformed into a 512 *×* 512 image (Eq. 16,17).

Sinusoidal dynamics at the flicker frequency (6 or 10 Hz) were applied to the images to interpolate them between the image and gray (Eqs. 13).

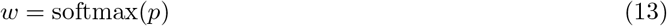

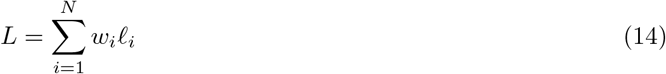

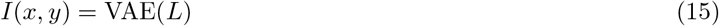

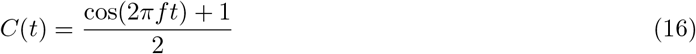

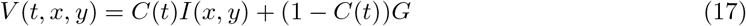

The resulting tensors were clipped to [0, 1], scaled by 255 and converted to integer values before being saved as mp4 files.

Latent textures 1 - 3 were created by sampling independent normal distributions with the VAE latent dimension (4, 64, 64). Latents 1 and 2 were spatially low-pass filtered. Latents 4 and 5 were black and white checkerboard patterns created in image space then passed through the VAE encoder. Latents 6 - 9 were images of natural objects taken by the first author and passed through the VAE encoder. Latent 10 was a white image passed through the VAE encoder.

Latents 11 - 40 were images taken from the Describable Textures Dataset and the Amsterdam Library of Textures [45, 46] cropped square and passed through the VAE encoder. A larger set of images were initially selected and these thirty images were sorted based on least similarity to previously added images computed from average color and distance in VAE latent space. The specific source files are listed in Section A.3.

### 6.6 EEG preprocessing and neural objective calculation

EEG was windowed to the 2 seconds preceding an LSL trigger which signaled the end of stimulation. All 68 EEG channels were referenced to the common average and 10 occipital and occipital-parietal channels were subselected for neural objective calculation, specifically Oz, O1, O2, PO7, PO3, POz, PO8, PO4, PO9, PO10. The same preprocessing was used for online optimization (MNE-LSL library [47] and offline analysis (MNE-python [48]).

On the 2-second preprocessed signal, a hann window was applied and the power spectral density was calculated for each channel. The bin closest to the flicker frequency was taken as the numerator and the mean of the bins *±* 2 bins away (1 Hz total) was used as the denominator. The log of this ratio was calculated per-channel and the mean across channels was used as the neural objective for optimization for each trial. In this paper, we use “log power spectral density ratio” (log PSD ratio) and “neural objective” interchangeably.

### 6.7 Optimizer and regression models

Bayesian optimization was used with a Bayesian Gaussian process as the surrogate model. Uniform priors were used for the squared-ex√ponential kernel lengthscale, GP variance, and diagonal noise variance, with distributions *U* (1e-3, *d*), *U* (1e-3, 1), and *U* (1e-2, 1), respectively. JAX [49] was used to construct the model and the NUTS sampler from numpyro [50] was used to draw samples from the posterior of the GP parameters.

To select the parameters for the next trial, the posterior samples were used to calculate the posterior predictive distribution, which was calculated per-sample for a set of 8 candidate parameter vectors. The upper confidence bound (UCB) was calculated for each sample from the posterior predictive distribution and the posterior-averaged UCB was calculated. L-BFGS-B [51] was used to optimize the candidate points with respect to the posterior-averaged UCB and the candidate with the highest UCB was selected as the candidate for the next trial. After each new datapoint was added to the optimizer, the neural objectives were re-zscored.

After the first block of preset trials was completed, the NUTS sampler was run for 250 warmup samples. Then, after each additional trial, 250 more steps were taken starting from the last sampler state and the 250 samples were decimated to 50 for use in Bayesian optimization.

### 6.8 Optimization effect size, ICC, confidence intervals, and ANOVA

For both optimization effect size (Hedges’ *g*) and intra-class correlation (ICC), we used the Pingouin Python library [52]. Hedges’ *g* is calculated as

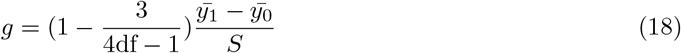

where *S* is the pooled standard deviation and df is the degrees of freedom in the standard deviation. For within-participant ICCs, the 2 repeats of each Preset stimulus are used and ICC(1,1) is reported [42]. For cross-participant ICCs each Preset trial is repeated both within and across participants. In this case, datasets were resampled 1000 times where one of the two within-participant repeats for each preset stimulus was randomly selected for each participant and the median “consistency” ICC(C, 1) was reported, which is invariant to each participant’s mean Preset neural objective. SciPy’s bootstrapping functions were used to calculate bootstrapped confidence intervals with the default bias-corrected and accelerated bootstrap confidence interval method [51]. The ANOVA was performed using the statsmodels Python library’s “ols” and “anova lm” functions.

### 6.9 Regression models

This same Bayesian Gaussian Process surrogate model was used offline as an encoding regression model. All settings were the same except for the number of warm-up samples (1000) and the number of posterior sampled generated (500 which were decimated by a factor of 10).

Ridge and RidgeCV models from scikit-learn [53] were also used for offline encoding regressions since it is straightforward to visualize the axis in predictor space that maximally modulates the predictions. For Ridge models trained on a single participant’s data, the “Leave-One-Out” cross validation strategy was used for the ridge parameter. For models trained on multiple participants’ data, cross-validation was performed leaving one training participant out to select the ridge parameter then the model was re-fit on all training data using the selected ridge parameter. We cross-validate a set of 41 ridge parameters log-spaced between 1 and 10^10^. For all regression models, we report linear correlations rather than the coefficient of determination since cross-validation of the Ridge parameter on neural datasets often leads to large regularization parameters which produces regimes where the model predictions track ground-truth data but at a much smaller absolute variance.

The video generator parameters were softmax transformed and then the square-root was taken (similar to a Hellinger distance [54]) before the parameters were zscored to the training mean and standard deviation for model fitting.

### 6.10 Image feature extraction

Image features extracted from the first frame (texture) of each video were:

- Luminance: mean of the Lab luminance channel across the image,
- Contrast: standard deviation of the Lab luminance channel across the image,
- Distance to gray: average RGB distance to gray across the image,
- R: average of the R channel across the image,
- G: average of the G channel across the image,
- B: average of the B channel across the image,
- Low-spatial frequency (SF): average log power of the lowest third of log-spaced frequencies (excluding 0),
- Mid-SF: average log power of the middle third of log-spaced frequencies, and
- High-SF: average log power of the highest third of log-spaced frequencies (up to the Nyquist frequency of 0.5 cycles/pixel for 512-pixel images).

The low-level image features were zscored to the training mean and standard deviation for model fitting.

### 6.11 Model weight distances

Pairwise model weight distances were calculated using the cosine distance

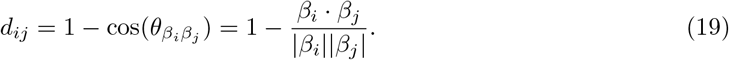

The mean cosine distances in Figure 12C,D are calculated in dimension *d* across *N* participants leading to 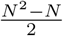 pairwise distances. The null distribution is estimated by drawing *N* random unit vectors in *d* dimensions, computing the pairwise cosine distances, and calculating the mean distance. This is repeated 50,000 times to generate the null distribution of means for calculating the p-values.

### 6.12 Texture modulation axis

For a fitted Ridge regression model, we want to define a trajectory through parameter space that is aligned with the model’s weight coefficient (modulation axis). Stimulus parameter changes parallel to this axis change the predicted neural objective, and stimulus parameter changes orthogonal to this axis leave it unchanged. However, the Ridge models are fit on transformed and zscored stimulus parameters, so we need to first define the trajectory in the linear model and then apply an inverse transform to the parameters. Let *x* and *y* be the transformed data and *x*_*z*_ and *y*_*z*_ be the transformed and zscored data. The fit Ridge model’s predictions look like

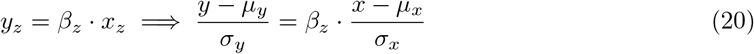

where the *µ* and *σ* variables are the mean and standard deviations of the training features. This can be rearranged to

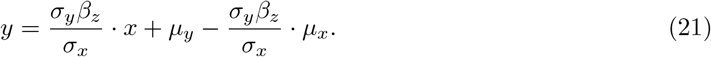

A linear trajectory can be defined by a point and direction. For the point, we use the mean *x* which leads to the mean *y* prediction *y* = *µ*_*y*_:

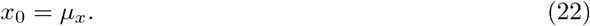

The unit-norm direction is the *β* for the unzscored parameters

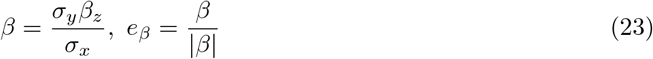

which means the linear trajectory in transformed parameter space is parametrized by

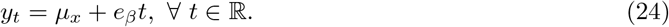

We set the limits of *t* for each experiment version by mean centering all transformed parameters used in the experiment and calculating their projections onto *e*_*β*_. The min and max projections set the limits for *t* and we take 11 linear spaced points along this trajectory for visualization.

To map back into untransformed stimulus parameter space, we square and then take the log of the transformed parameters. These vectors are passed into the video generator and the first frame of each is shown in Figure 12E.

## 7 Discussion

In this work, we describe a system for closed-loop optimization that combines a parameterized generative image model latent search space and flicker dynamics to modulate steady-state visual evoked potentials (SSVEPs) as recorded by EEG in the theta and alpha frequency bands. The optimization of SSVEP response can be performed reliably in 10, 20, and 40 dimensional parameter spaces, although the resulting optimization effect size is smaller for higher dimensions. Stimuli and encoding model predictions derived from optimization participants generalize to new participants not in the training set. The distances between regression model weights from individual participants are closer than what would be expected by chance, revealing shared visual neural response dependence on stimulus parameters across individuals. Together, our results show that closed-loop optimization of stimulus parameters is a viable tool for studying rhythmic entrainment of visual responses using EEG.

Recent evidence from both neurophysiological and modeling studies suggests that rhythmic neural activities observed across various recording modalities are influenced by both endogenous oscillatory dynamics and their entrainment, or coupling, to periodic external stimulation (see review by [55]). Specifically, alpha-band SSVEP has been shown to be not a mere superposition of a series of evoked potentials, but entrainment of oscillations detectable by both noninvasive and invasive methods [56, 57]. Modulation of alpha oscillation with flashing lights has also been shown to modulate visual perception in human participants [58]. Moreover, the optimization of SSVEP in alpha and theta frequencies result in different weight profiles of visual textures and features, suggesting the neural response function to some visual features vary between the two frequencies (Fig. 12B). Since the current study focuses on the validation and characterization of the closed-loop system, we did not try to further disentangle the interaction between endogenous and stimulus-driven activities. This may be achieved via closed-loop manipulation of SSVEP phase instead of amplitude, because stimulus-driven activities are phase-locked to the visual input in a predictable way while the endogenous activities are not.

Beyond theta and alpha, we expect that the same paradigm could be used to develop optimized stimuli for entraining SSVEP oscillations at any frequency supported by the display system. This would enable clinical researchers to explore non-invasive, optimized stimulation techniques in diseases where modulating endogenous neural oscillations through entrainment may have therapeutic benefit. The closed-loop optimization pipeline is not constrained to sensory stimulation and EEG, but can be adapted to other stimulation methods including transcranial electric and magnetic stimulation methods such as TMS and tACS. Compared to transcranial methods, visual (sensory) modulation does not directly apply potentiation or inhibition to local circuits [59, 60]. Nevertheless, the rich parameter space of sensory stimuli allows optimization for a broader range of neural objectives, including non-periodic, time-varying ones and objectives in functional areas downstream of early visual cortex.

Outside of the flicker and SSVEP paradigm, however, there may not be clear choices for what spatiotemporal stimulus features will evoke neural objectives of interest and optimization may be more challenging if there is only a weak relationship between stimulus parameters and neural objective modulation. To improve optimization performance for higher dimensions in this case, large scale (open) data collection can be used to train encoding models [61, 62, 63, 64] that can move the shoulder in *P* (*d*) in Figure 10B up and to the left since the neural responses as a function of stimulus parameters generalize across participants (Fig. 11). Similarly, optimizing neural objectives outside of visual cortex may be comparatively more difficult using visual stimulation, but a multimodal stimulus space and strong encoding model priors could flexibly target a wider range of neural objectives across diverse functional areas. These improvements to search efficiency may also allow closed-loop systems to scale to stimulus search spaces much larger than those considered here such as the full variational autoencoder latent space (∼16,000d) or a CLIP vector space (∼1,000d)[65] which could be used in multimodal stimulation, for example video, audio, and text.

More broadly, we view our contributions as a step toward a closed-loop system which could modulate a wide range of neural targets using a diverse set of sensory-, electromagnetic-, and implant-based stimulus parameter spaces. Such a system which would have enormous benefit for basic neuroscience and clinical neuromodulation applications.

## 8 Author contributions

J.A.L., Y.S., S.W., D.J.K., and A.H. conceived and designed and work. J.A.L. and S.W. collected the data., J.A.L., Y.S., and A.I. wrote the analysis code and performed the analysis. J.A.L. and Y.S. wrote the paper.

## Appendix A Extended Data

### A.1 Participant-experiment version mapping

#### A.2 ANOVA

Table 1 shows the summary of the ordinary least-squares fit underlying the ANOVA and Table 2 shows the sums of squares.

**Table 1:**
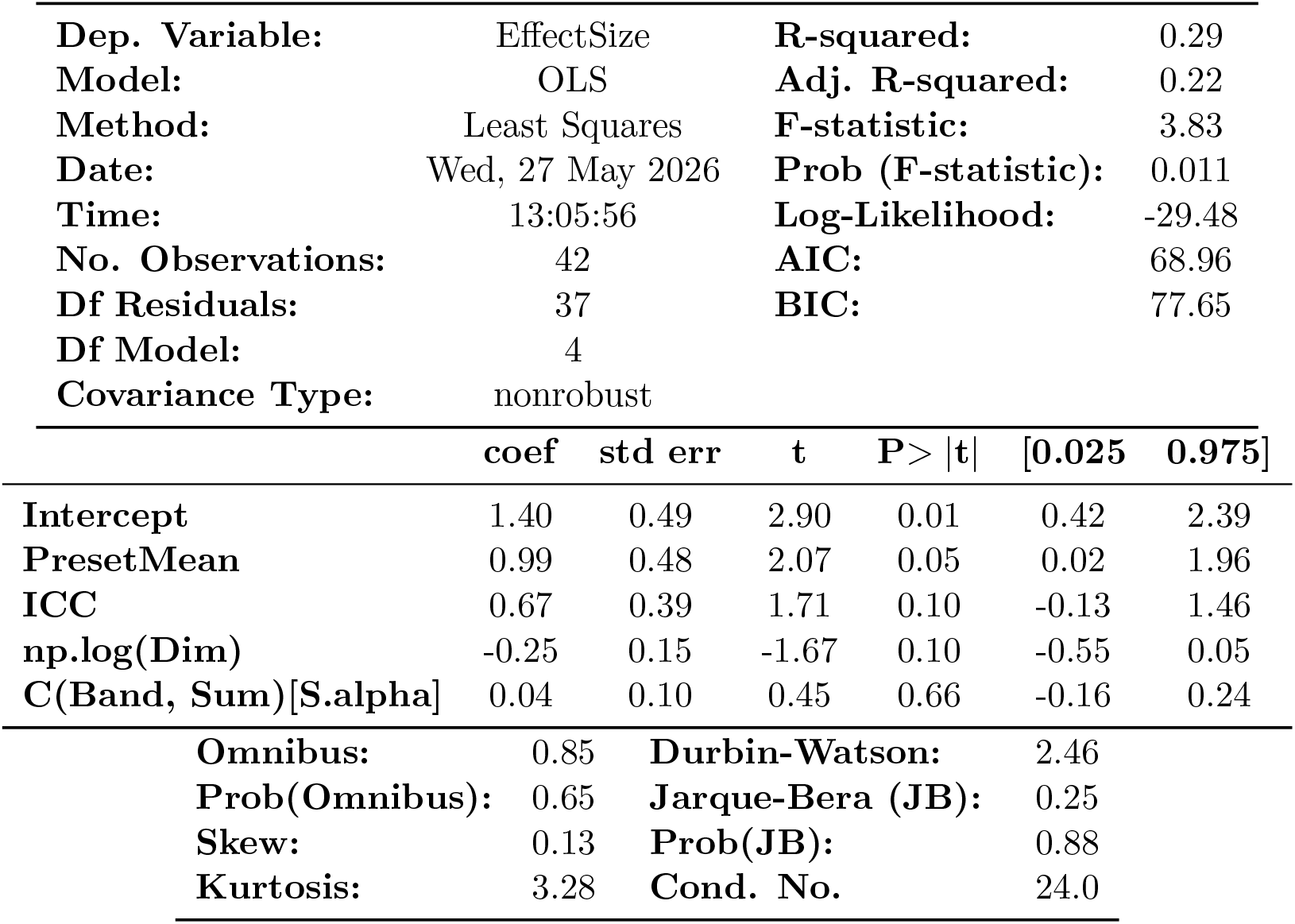
OLS Regression Results.

|  |  |  |  |
| --- | --- | --- | --- |
| **Dep. Variable:** | EffectSize | **R-squared:** | 0.29 |
| **Model:** | OLS | **Adj. R-squared:** | 0.22 |
| **Method:** | Least Squares | **F-statistic:** | 3.83 |
| **Date:** | Wed, 27 May 2026 | **Prob (F-statistic):** | 0.011 |
| **Time:** | 13:05:56 | **Log-Likelihood:** | -29.48 |
| **No. Observations:** | 42 | **AIC:** | 68.96 |
| **Df Residuals:** | 37 | **BIC:** | 77.65 |
| **Df Model:** | 4 |  |  |
| **Covariance Type:** | nonrobust |  |  |

|  | coef | std err | t | P>\|t\| | [0.025 | 0.975] |
| --- | --- | --- | --- | --- | --- | --- |
| **Intercept** | 1.40 | 0.49 | 2.90 | 0.01 | 0.42 | 2.39 |
| **PresetMean** | 0.99 | 0.48 | 2.07 | 0.05 | 0.02 | 1.96 |
| **ICC** | 0.67 | 0.39 | 1.71 | 0.10 | -0.13 | 1.46 |
| **np.log(Dim)** | -0.25 | 0.15 | -1.67 | 0.10 | -0.55 | 0.05 |
| **C(Band, Sum)[S.alpha]** | 0.04 | 0.10 | 0.45 | 0.66 | -0.16 | 0.24 |
| **Omnibus:** | 0.85 | **Durbin-Watson:** | 2.46 |
| **Prob(Omnibus):** | 0.65 | **Jarque-Bera (JB):** | 0.25 |
| **Skew:** | 0.13 | **Prob(JB):** | 0.88 |
| **Kurtosis:** | 3.28 | **Cond. No.** | 24.0 |

**Table 2:** ANOVA Results.

|  | sum_sq | df | F | PR(>F) |
| --- | --- | --- | --- | --- |
| PresetMean | 1.16 | 1 | 4.29 | 0.05 |
| ICC | 0.79 | 1 | 2.91 | 0.10 |
| np.log(Dim) | 0.76 | 1 | 2.80 | 0.10 |
| C(Band, Sum) | 0.05 | 1 | 0.20 | 0.66 |
| Residual | 10.01 | 37 | NaN | NaN |

#### A.3

Images from the Describable Textures Dataset: bumpy 0059.jpg, grid 0025.jpg, meshed 0031.jpg, paisley 0023.jpg, polka-dotted 0069.jpg, scaly 0113.jpg, zigzagged 0047.jpg.

Images from the Amsterdam Library of Textures: 108 c1l1r120.png, 140 c1l1r120.png, 145 c1l1r120.png, 14 c1l2r60.png, 174 c1l1r60.png, 176 c1l1r180.png, 180 c1l1r180.png, 182 c1l1r120.png, 197 c1l1r60.png, 213 c1l1r120.png, 216 c1l1r60.png, 225 c1l1r60.png, 23 c1l2r120.png, 27 c1l2r60.png, 34 c1l3r60.png, 39 c1l2r120.png, 45 c1l1r120.png, 47 c1l1r180.png, 48 c1l1r180.png, 59 c1l1r180.png, 5 c1l3.png, 79 c1l1r120.png, 80 c1l1r120.png.

